# Physical theory of biological noise buffering by multi-component phase separation

**DOI:** 10.1101/2021.01.05.425486

**Authors:** Dan Deviri, Samuel A. Safran

## Abstract

Maintaining homeostasis is a fundamental characteristic of living systems. In cells, this is contributed to by assembly of biochemically-distinct organelles, many of whom are not membrane-bound, but form by the physical process of liquid-liquid phase separation (LLPS). By analogy with LLPS in binary solutions, cellular LLPS was hypothesized to contribute to homeostasis by facilitating “concentration buffering”, which renders the local protein concentration within the organelle robust to global variations in the average, cellular concentration (e.g. due to expression noise). Interestingly, concentration buffering was experimentally measured in vivo, in a simple organelle with a single solute, while it was observed not to be obeyed in one with several solutes. Here, we formulate theoretically and solve analytically a physical model of LLPS in a ternary solution of two solutes (A and B) that interact both homotypically (A-A attractions) and heterotypically (A-B attractions). Our physical theory predicts how the equilibrium concentrations in LLPS are related to expression noise and thus generalizes the concept of concentration buffering to multi-component systems. This allows us to reconcile the seemingly contradictory experimental observations. Furthermore, we predict that incremental changes of the homotypic and heterotypic interactions among the molecules that undergo LLPS, such as those that are caused by mutations in the genes encoding the proteins, may increase the efficiency of concentration buffering of a given system. Thus, we hypothesize that evolution may optimize concentration buffering as an efficient mechanism to maintain LLPS homeostasis, and suggest experimental approaches to test this in different systems.

**Significance:** The stochastic nature of transcription/translation implies that the concentrations of cellular proteins are “noisy” and not constant in time or across cell populations. Liquid-liquid phase separation (LLPS), can reduce or “buffer” this noise by maintaining well-defined concentrations, even in the presence of concentration distributions. However, this idea was recently challenged experimentally in multicomponent systems. Our physical theory of LLPS in ternary systems (solutes A and B in a solvent), predicts their LLPS properties as a function of the A-A (homotypic) and A-B (heterotypic) interaction strengths. We show how buffering can be effective if the noise distribution aligns with the tie-lines of the phase diagram and suggest that evolution may optimize concentration buffering by selecting mutations that lead to this.

## 1 Introduction

Over the past decade, a significant body of research has revealed the crucial involvement of liquid-liquid phase separation (LLPS) in the formation of many membraneless organelles, also known as bio-molecular condensates (BMCs) [1], among which are the nucleolus [2], P-granules [3], stress-granules [4], and heterochromatin [5]. The distinct biochemical environment of the BMCs (e.g. enzymes, substrates) can promote reactions related to important biological functions. This property is similar to membrane-bound organelles. However, in contrast to membrane-bound organelles, whose homeostasis is maintained by the out-of-equilibrium action of active pumps and channels embedded in the membrane, the concentration homeostasis in BMCs is established via thermodynamic equilibrium due to attractive interactions among molecules and the solution entropy of the phase separating components. In several situations, such as those of the different types of ribonucleoprotein bodies [6], the BMCs are formed in systems with two or more solutes. For the situation of two solutes with interactions that dominate the mixing tendency of the entropy (proportional to the temperature), one can think of two simple limits: Attraction between molecules of one species (homotypic) leads to segregative LLPS where the other solute is mostly excluded from the BMCs, while attraction between molecules of different types (heterotypic) lead to associative LLPS where both solutes cooperatively form BMCs (see Fig. 1) [7]. In the more realistic case, combinations of homotypic and heterotypic attractions, can drive diverse types of LLPS, leading to a rich variety of BMCs, that may themselves consist of multiple phases, such as the layered nucleolus [2]. In this work, if not specified otherwise we consider LLPS into two phases, which gives rise to a one phase which is a BMC that thermodynamically coexists with a phase that is (much) more dilute in the solute concentrations.

**Figure 1:**
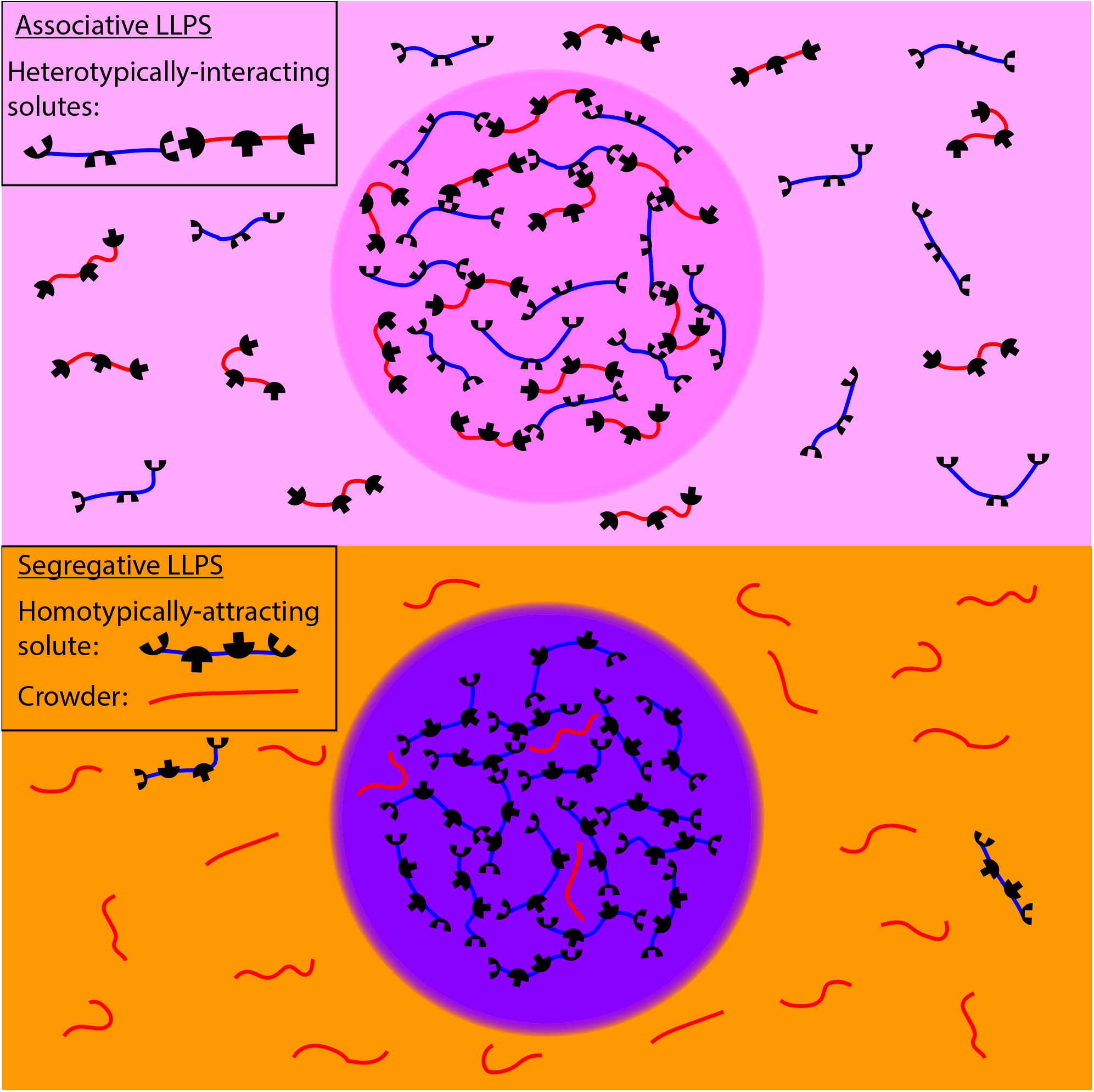
Illustration of associative vs. segregative LLPS. Top: Two types of solute molecules (red and blue) that interact heterotypically (attractions of blue and red) undergo associative LLPS and form a dense phase rich in both solutes (light purple), that coexists with a solution dilute in both solutes (light pink). Bottom: A homotypically-attractive solute (where blue attracts blue) and an inert “crowder” (red) undergo segregative LLPS and form a phase rich in the attractive solute but poor in the crowder (purple), and a phase rich in crowder but poor in attractive solute (light orange).

The large number of BMCs and apparent ubiquity of LLPS as an organizational principle of organelles in living cells suggests that it confers some evolutionary advantages. One of the hypothesized advantages is mitigating the deleterious outcomes of “expression noise”. Cells, even when carrying identical genomes, exhibit natural variations of the concentrations of transcription and translation related molecules, such as transcription factors, polymerases, or ribosomes. This variation introduces cell-to-cell fluctuations in the production and subsequent concentrations of cellular molecules, which, when measured in an ensemble of genetically identical cells, follow a statistical distribution rather than deterministic values [8]. For biochemical processes that require a narrow concentration range of molecules to be faithfully executed, expression noise can be detrimental.

LLPS may mitigate the harmful effects of expression noise by facilitating “concentration buffering”: Fluctuations in the overall (spatially-averaged) concentration of the molecules are “buffered” (shielded), so that all the changes in the overall concentration are merely absorbed in changes of the volumes of the phases, while the concentrations of the molecules within each phase remain unchanged at fixed equilibrium values. This allows the biological function of the BMCs, which may be sensitive to the concentrations, to remain robust to their stochastic fluctuations [1, 9, 10]. This “simplistic” type of concentration buffering, exists only in LLPS of binary solutions, consisting of a single solute in a solvent; in the multi-component case, applicable to the cellular environment, changes of protein concentrations within different phases were experimentally observed as the overall protein concentrations were varied [11, 12, 13, 14]. At first glance, this shows the lack of concentration buffering in the multi-component system. However, in this paper, we theoretically demonstrate how concentration buffering is nevertheless maintained in multi-component solutions, with a specific example of the case of two solutes in a solvent which can form either segregative or associative phase-separated volumes. The buffering is more subtle and constrained than in the simplistic case, but can be nonetheless be effective, depending on the correlations of the noise in the expression of the two components as we predict.

In binary solutions, the interplay between molecular attraction, which promote LLPS, and temperature, which inhibits LLPS (by increasing the role of entropy via molecular collisions and mixing) determines the conditions for which a solution can phase separate in equilibrium and the subsequent concentrations of the two coexisting phases. Those concentrations are independent of the overall (system averaged) solute concentrations, which fix only the relative volumes of the two phases [15]. This independence of the coexisting concentrations on the overall concentration, is the physical basis for concentration buffering in binary solutions. Recent experimental study [16] have demonstrated that even in the out-of-equilibrium, multi-component cellular environment, concentration buffering may decrease local concentration fluctuations. This will be the case when the timescales for protein diffusion that establishes the spatial separation of the two phases, is shorter than those of gene expression and degradation [16], responsible for the concentration variations. Therefore, on these shorter time scales, the phase separation is established in a quasi-static equilibrium manner. While it is clear why buffering may be effective in the out-of-equilibrium conditions of the cell, it is unclear how buffering is impacted by its multi-component nature, in which the equilibrium concentrations of molecules within the coexisting phases generally depend on their overall concentration [17].

As we explain in section “Concentration buffering in solutions of two solutes” below, despite this general dependence in multi-component LLPS, there is always a special, correlated change of the overall concentrations that nevertheless does not change the coexisting concentrations; this special direction is given by the tie-lines on the phase diagram as function of the two concentrations. Tie-lines connect the two coexisting phases [18], and a system prepared with an average concentration along the tie-line always separates into the same two coexisting phases; the concentration changes along the tie-lines only modify the relative volumes of the two phases. If the expression noise of multiple genes which encode the molecular components of phase separating system is correlated by the underlying gene network in a manner that matches the tie-line, its effect on the coexisting concentrations is buffered. We hypothesize that, for the biological contexts in which expression noise is deleterious, alignment of the correlated expression noise with the tie-line may increase the fitness of the organism since such alignment means that the compositions of the two coexisting phases will be (approximately) unaffected by the noise. Thus, evolution might fine-tune the correlation of the gene expression noise, the tie-lines of the phase diagram, or both to mitigate the harmful effects of expression noise on LLPS-based concentration buffering. We elaborate upon this in a pictorial manner in the next section, and quantify and classify in a generic manner the relevant phase diagrams in the section on Results.

This is preceded by a section that introduces our model which treats in a unified manner LLPS driven by both homotypic and heterotypic interactions. In the results section, we plot the phase diagram and predict the tie-lines for three archetypical cases, all of which are included in our generic model: (1) A solution of a scaffold (self-attracting), crowder (inert), and a solvent that undergo segregative LLPS with characteristic, negatively-sloped tie-lines. (2) A solution of two mutually attracting molecules and a solvent that undergo associative LLPS, with characteristic, positively-sloped tie-lines. (3) A solution of a scaffold, a client (a binding partner of the scaffolds) and a solvent, which we find may undergo segregative or associative LLPS with tie-lines of variable slopes, depending on the relative interaction parameters.

In the discussion section, we use our predictions of the directions of the tie-lines for these three archetypical solutions to delineate scenarios in which there may be possible alignment between the direction of the expression noise and the tie-lines. There are also other situations where the nature of the tie-lines can still shield BMCs from biochemical changes in the environment even without complete alignment. Based on the regulatory network associated with the proteins involved, the outer layer of the nucleolus may be an example of alignment of the noise and the tie-lines. We suggest future experiments required to further test our evolutionary hypothesis and discuss the possible evolutionary mechanisms of alignment. Surprisingly, our model predicts that under certain conditions, archetypical solutions of multivalent scaffold and multivalent binding partner can exhibit re-entrant LLPS as the heterotypic interaction strength is varied, a prediction that is qualitatively supported by recent experimental observations [19]. In addition, we discuss how concentration buffering can be generalized from a three-component to a system with a larger number of components that phase separates into multiple phases, and predict that in such cases the buffering becomes more effective as the number of equilibrium phases increases.

## 2 Concentration buffering in solutions of two solutes

In contrast to the fixed, coexisting concentrations in LLPS of a single solute that gives rise to the simplistic type of concentration buffering, in LLPS of solutions with two (or more) solutes, the two coexisting concentrations of solutes do vary with the overall concentrations [11]. Nevertheless, there is a direction in the phase diagram (as a function of the concentrations) of the two solutes, along which concentration buffering can be defined. This direction is that of the tie-lines in the concentration-concentration phase diagrams at fixed temperature and interaction energies. The variation of overall concentration along the tie-lines always results in coexisting phases with same concentrations (buffering); the changes in concentration only modify the relative volumes of the two coexisting phases. Therefore, the two ends of the tie-line, specify the concentrations of the two coexisting phases. However, variations of the overall concentrations that deviate from those along the tie-line generally bring the system to an equilibrium defined by other tie-lines that lead to different coexisting concentrations (see Fig. 2A). Thus, in the multi-component, cellular environment, only correlated variations of the overall (system average) molecular concentrations that align with the directions of the tie-line will be “buffered”. We now outline the cases for which expression noise of multiple genes is correlated and may be aligned with the tie-lines and buffered.

**Figure 2:**
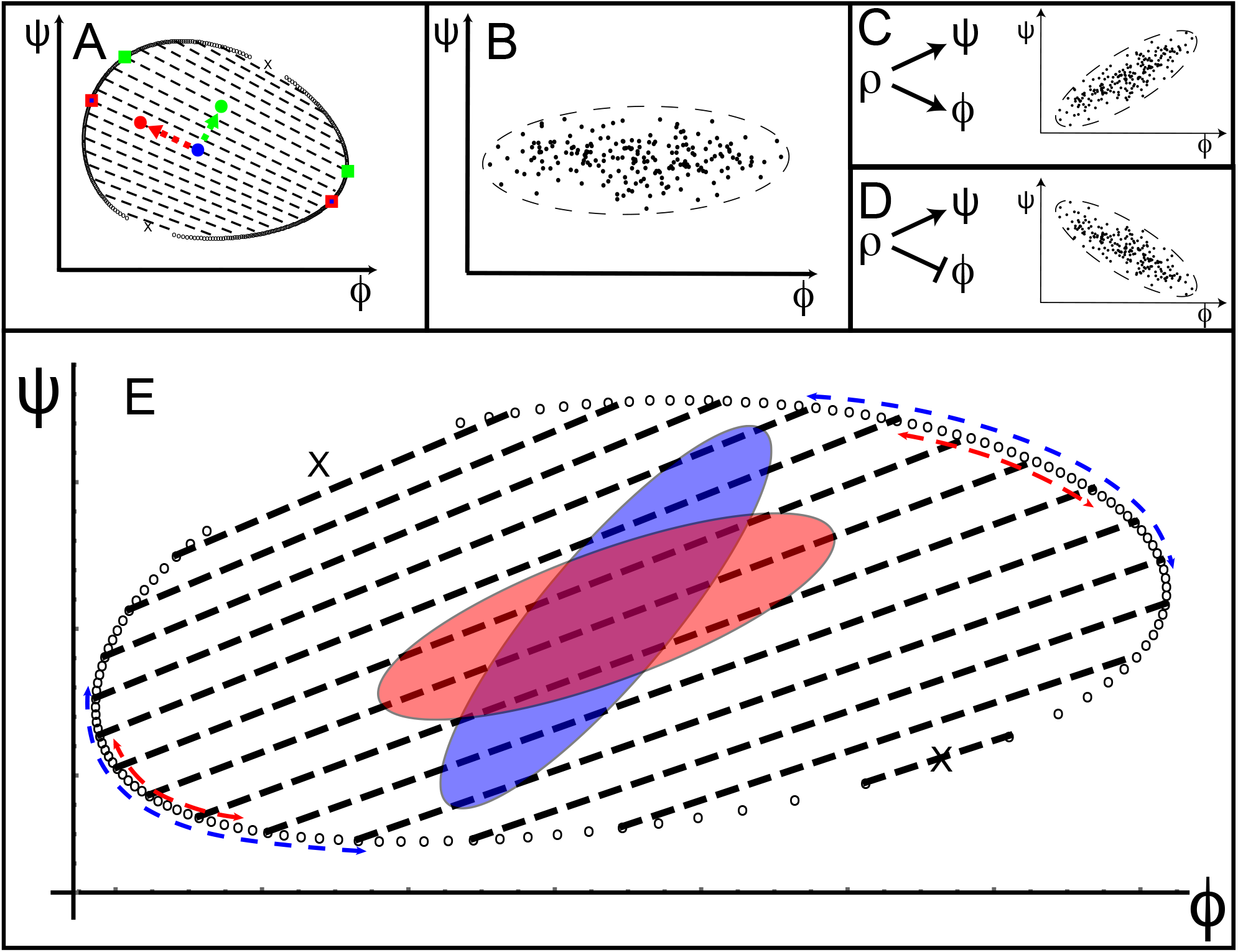
Biological noise and concentration buffering. (A) Consequences of variations in the average (overall) concentrations on the concentrations of each of the solutes in the two coexisting phases (full squares) along and perpendicular to the tie-lines (black dashed lines); the ties-lines end at the concentrations of the two coexisting phases (open black circles). A variation of the initial concentrations of the two solvents, represented by the blue circle, along the direction of the red arrow, that is aligned with the tie-lines, does not change the coexisting concentrations (red square with blue dot in center). In contrast, a variation of the same initial concentrations in the direction of the green arrow perpendicular to the tie-lines, changes the coexisting concentrations, which now have the values indicated by the green squares. (B) Significant part of the distribution of the overall concentrations and its envelope, the “noise-ellipse”, when only intrinsic noise is present. The intrinsic noise in the expression of solute *ϕ* is more prominent, so that the major axis of the noise-ellipse is parallel to the *ϕ* axis. (C and D) Tilted noise-ellipses in the presence of both intrinsic and extrinsic noises, the latter originates from concentration fluctuations of a common transcription factor *ρ*. In (C), *ρ* induces expression of both *ϕ* and *ψ*, while in (D) it induces *ψ* and represses *ϕ*. (E) The relation between the alignment of the tie-lines (dashed black lines) and the orientations of the noise-ellipses (blue and red ellipses), determines the degree of concentration buffering. The axes of the two noise-ellipses are equal in lengths, but the direction of the red ellipse is aligned with the direction of the tie-lines while the blue ellipse is not. The variation in the concentrations of the two solutes in the coexisting phases (dashed blue arrows) due to the blue noise-ellipse, is significantly larger than the variation in the equilibrium concentrations (marked by the red arrows) due to the red noise-ellipse.

If there were no expression noise, the concentration of proteins in genetically identical cells would be the same, and their steady state levels would be constant in time. In that case, the overall concentrations of proteins of an ensemble of cells would be described by a single point in the phase diagram. However, biological (and thermal) noise is always present, so that even genetically identical cells differ with respect to their protein abundance, for example due to the stochastic nature of the transcription and translation processes of each gene independently [20]. These contributions to the noise are “intrinsic” to the expression of each gene, so that the noises in the expression of different genes are uncorrelated. Due to the intrinsic noise, the overall concentrations of proteins in an ensemble of cells are thus described by a distribution of points in the phase diagram, centered around the average concentrations of each protein, rather than a single point. The shape of the distribution due to the intrinsic noise only depends on the relative magnitudes of the expression noise of the two genes. If the magnitudes of the two expression noises are equal, then the distribution is a circle centered at the average values; however, if the expression noise of one gene is larger than the other, then the distribution is an ellipse whose major axis is parallel to the concentration axis of the gene which is more susceptible to the noise (see Fig. 2B). The envelope of the distribution, defined as the locus of points for which this distribution is significant (e.g, the contour that enclose 90% of the points) characterizes the “noise-ellipse”, in which variations of the two concentrations are significant.

In addition to the intrinsic factors, there are “extrinsic” factors that affect the expression of different genes in a correlated manner [21]. For example, these factors can be variations of the concentrations of “master” proteins that are involved in the expressions of multiple genes, such as RNA polymerases, ribosomes, or transcription factors. The concentration variations of these master proteins may originate from intrinsic noise in the expression of their own encoding genes, but their effect on the expression of the downstream genes, in which expression they are involved, is extrinsic. Because the extrinsic noise can simultaneously affect the expression of multiple genes, its contributions to the variations in the concentrations of the expressed proteins can be correlated. These correlations affect the major and minor axes of the noise-ellipse but more importantly, rotate the axes of the noise-ellipse with respect to the axes of the solute protein concentrations. Therefore, the sum of both the extrinsic and intrinsic noises results in a concentration distribution (of the two genes in the phase diagram) which is a tilted ellipse (see figures 2C and 2D).

A simple case is one in which the extrinsic noise is dominated by concentrations fluctuations of a single protein that regulates the expression of multiple target genes (e.g. a transcription factor [22]). In that case, the lengths of the axes of the noise-ellipse and its orientation can be analytically expressed as functions of the magnitude of the noises and biochemical properties of the proteins (degradation rates and their susceptibilities to the noise of the regulator protein) [23]. If the intrinsic noises are similar in magnitude, the orientation of the noise-ellipse is primarily determined by the protein degradation rates and the susceptibilities of their promoters to the transcription factor, but is independent of the magnitude of the intrinsic and extrinsic noises. For example, the concentrations of two proteins whose expression is induced by the same transcription factor is positively correlated, but is negatively correlated if the same transcription factor induces the expression of one, but represses the other (see figures 2C and 2D).

The biological effect of this expression noise is, in some cases, beneficial, however in others it is deleterious and must be mitigated. One way this can occur is when the major axis of the noise-ellipse is aligned with the tie-lines of the phase diagram. In that case, most of the noise shifts the two concentrations along the tie-lines and the resulting equilibrium concentrations of the two coexisting phases remain fixed (i.e., show concentration buffering). Of course, the noise in the direction of the minor axis, does shift the system from one tie-line to another, but if this axis is relatively small compared to the major axis, the shift will be insignificant and the two coexisting phases will be almost the same as in the ideal, noiseless case. As shown in Fig. 5E, concentration buffering may thus reduce the noise in a multicomponent solution, with increasing efficiency as the orientations of the noise-ellipse and the tie-lines of the phase diagram are more aligned. That indeed may be a fitness criterion which evolution will cause to be satisfied. The orientation of the tie-lines depends on several generic features of the intermolecular interactions. We present a model that allows us to predict the phase diagrams and tie-line orientations for three archetypical cases in the following section.

## 3 Theoretical model of multi-component LLPS

In this section, we present a physio-chemical model that predicts the phase diagrams and tie-lines for systems that comprise two solutes in a solvent. This allows us to predict the generic features of the phase diagrams that are relevant to the problem of concentration buffering. The free energy of our model, which is a generalization of the Flory Huggins (FH) model for binary polymer solution, takes into account both homotypic and heterotypic interaction and allows us to investigate their distinct effects on the LLPS properties of the solution in the following results section. The generalization of this approach to systems with more than two solutes is discussed in the SI.

The two solutes interact via short-range interactions (e.g. proteins in the cytoplasm), and we write an approximate, mean-field expression for the free energy that includes both the entropy of mixing and interaction energies. The free energy per unit volume *f* (*ϕ, ψ*), is a function of the local volume fractions of the two solutes *ϕ, ψ*; the volume fraction of solvent *ϕ*_*s*_ (for a system of incompressible molecules) is then fixed at *ϕ*_*s*_ = 1 −(*ϕ* + *ψ*).

The function *f* (*ϕ, ψ*) contains all the information needed to characterize the equilibrium state of the system, which minimizes the total free energy of the system, subject to the constraint that the total number of molecules of each type are conserved. These imply [15] that in equilibrium, the free energy change resulting from the addition or removal of a molecule of any type from each phase, known as its chemical potential, must be the same in each of the coexisting phases. If the chemical potentials for the different phases are different, the free energy is not minimal because a non-equilibrium flux of molecules from a phase of high chemical potential to a phase of low chemical potential, decreases the total free energy. Second, the equilibrium free energy change resulting from an increase or decrease of the volume of each of the coexisting phase, known as its osmotic pressure, must also be the same in each phase. Similar to the case of chemical potentials, an imbalance of the osmotic pressures, results in a non-equilibrium expansion of the high-pressure phase at the expense of the low-pressure phase (i.e., resulting from a flux of solvent molecules); this would decrease the total free energy of the system, indicating that the total free energy is not minimal. The conditions of equality of chemical potentials and the osmotic pressures are expressed, in terms of the free energy per unit volume *f*, as the following equations for two-phase equilibria (see SI):

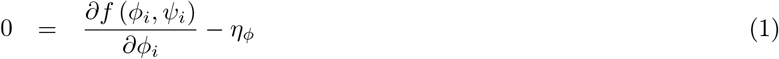

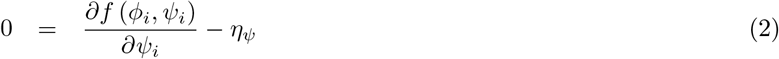

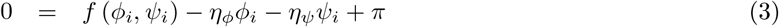

where *ϕ*_*i*_ and *ψ*_*i*_ are respectively the concentrations (in terms of volume fractions) of the solute molecules *ϕ* and *ψ* in phase *i*, where *i* = 1, 2, *η*_*ϕ*_ and *η*_*ψ*_ are respectively the chemical potentials of solutes *ϕ* and *ψ*, and *π* is the osmotic pressure of the phases. In the SI, we generalize this to systems with more than two solutes and two equilibrium phases, and also show how the thermodynamic equilibrium conditions dictate the volumes and numbers of molecules of each type in the coexisting phases.

To predict the phase diagrams of biological systems, we must specialize the model to the situation of LLPS of biomolecules. In biological systems, the majority of the phase separating molecules are multivalent intrinsically disordered proteins (IDPs) and RNA molecules [24]. These molecules are usually described as a collection of different “stickers” connected by “spacers” (e.g. polymeric backbone), where a group of stickers of specific types may interact strongly [24]. Homotypic interactions refer to stickers belonging to the same type of molecules., while heterotypic interactions refer to stickers belonging to molecules of different types. Molecules that contain different kind of stickers may potentially present a combination of homotypic and heterotypic interactions. In solutions, most IDPs and RNA molecules usually appear as open, unstructured coils rather than compact globules [25, 26]. Therefore, stickers and spacers of different molecules are accessible to each other and may interact. As an approximation, we follow the FH model in which all of the monomeric segments uniformly contribute to the interactions, regardless of their position along the polymer backbone. This is different than compact globules, such as folded proteins, whose surface amino acids are the only ones that participate in inter-molecular interactions, in which case the mean-field treatment of FH theory may be inappropriate [27].

For a ternary solution comprising two solutes in a solvent, the free energy density *f* in the FH theory is:

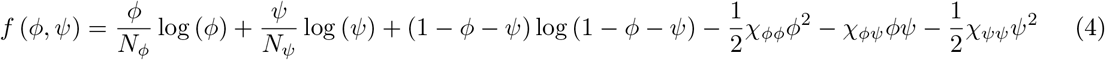

where *ϕ* and *ψ* are the concentrations of the two polymers, *N*_*ϕ*_ and *N*_*ψ*_ are the number of persistence lengths [28] of each polymer and the set of {*χ*_*ij*_} are the FH interaction parameters, with *χ*_*ϕϕ*_ and *χ*_*ψψ*_ accounting for the homotypic, (self-attractive) interaction energies of monomers of type *ϕ* and *ψ* respectively, while *χ*_*ϕψ*_ is the heterotypic interaction energy related to interaction between the monomers of the two different types, *ϕ* and *ψ*. We use the convention in which all the interaction energies are divided by the thermal energy *k*_*B*_*T*, and are hence dimensionless.

We note that FH models with a single interaction parameter for every pair of molecule types, which averages the relevant stickers over the entire polymer, has been shown to approximate well the LLPS properties of heterogeneous stickers-and-spacers proteins, both in experimental [11] and theoretical [29] contexts. We next use Eqs. 1, 2, and 3 to plot the phase diagrams and tie-lines for concrete examples of biological interest.

## 4 Results

Most BMCs consist of a single phase with a relatively high concentration of one or more proteins which coexists with an aqueous phase where the proteins are dilute. This coexistence results from LLPS that forms two phases. Furthermore, in BMCs, usually there is no more than a single multivalent scaffold protein that can phase separate from the solvent in a binary solution. Motivated by these observations, we focus on ternary solutions in which one of the solutes, *ψ*, has no homotypic attraction, by setting *χ*_*ψψ*_ = 0. Indeed, as detailed in the subsections below, we find that such ternary solutions can phase separate into only *two* phases. In the SI, we discuss the scenario of LLPS into more than two phases, which may provide an even more robust concentration buffering.

When *χ*_*ψψ*_ = 0, there are three archetypical combinations of the homotypic interaction parameter *χ*_*ϕϕ*_ and the heterotypic one *χ*_*ϕψ*_: (1) *χ*_*ϕϕ*_ > 0 and *χ*_*ϕψ*_ = 0, in which *ϕ* can phase separate into a phase with a relatively high concentration of *ϕ* that coexists with an aqueous phase that also contains *ψ*. (2) *χ*_*ϕϕ*_ = 0 and *χ*_*ϕψ*_ > 0, in which neither *ϕ* nor *ψ* separately phase separates into a relatively high-concentration phase, but can phase separate into a phase that contains a relatively high-concentration of them both, due to the strong, heterotypic interactions. (3) *χ*_*ϕϕ*_ > 0 and *χ*_*ϕψ*_ > 0, which combines the two former cases. In this scenario, the presence of both homotypic and heterotypic interactions can either drive LLPS with relatively high concentrations of both molecules or a phase with a relatively high concentration phase of only one of them, depending on the relative strengths of the two interactions. Each of these cases is analyzed below, and the appropriate phase diagrams are plotted using Eq. 4 for the free energy with the relevant interaction parameters. Eqs. 1, 2, and 3 are then used to predict the binodal and tie-lines. To find the critical points (marked by “X”), which are points in concentration-space in which the compositions of the two coexisting phases are identical, we followed the method presented in third section of the SI of [30].

### 4.1 Case I – Homotypic (*χ*_*ϕϕ*_ > 0, *χ*_*ϕψ*_ = 0): Concentrated phase rich in one solute (segregative LLPS)

Biological environments such as the cytoplasm contain a large number of macromolecules that do not significantly or specifically interact with proteins that undergo homotypic LLPS, but of course, cannot occupy the same location in space as the other proteins or solvent molecules (excluded volume interactions); such macromolecules are referred to as crowders. Excluded volume interactions introduced by crowders are inherently accounted for in FH theory by its construction. Thus, we model crowding, by considering one solute *ϕ* with homotypic (self-attractive) interactions *χ*_*ϕϕ*_ > 0 and another, *ψ*, that interacts attractively neither with *ϕ* nor itself (*χ*_*ϕψ*_ = *χ*_*ψψ*_ = 0).

In Fig. 3, we plot two concentration-concentration phase diagrams for this case, calculated using typical, homotypic interaction strengths. We identify three classes of phase diagrams: The first type, presented in Fig. 3A, is the one in which the binodal intersects the *ψ* = 0 axis, indicating that the solute *ϕ* can phase separate even in the absence of crowder solutes. The quantitative ranges of the molecular interactions and sizes for which this occurs is determined by the condition 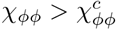, whose dependence on the interactions and molecular lengths is given in the SI, as it is for all the other critical values of interaction parameters; the characteristic values of the interaction parameters and the topological properties of the phase diagrams that are associated with them are listed in Tables 1 and 2 in the SI. The second type, presented in Fig. 3B, is the one in which the binodal does not intersect the *ψ* = 0 axis, indicating that crowding is required for LLPS of solute *ϕ*. This type is characterized by interaction energies in the range 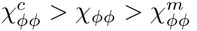 (see SI). The third type of phase diagram, which we do not plot, is one in which LLPS does not occur and the single phase, homogenous solution is the equilibrium state for any concentrations *ϕ, ψ* (see SI). This happens when the homotypic interaction is smaller than another critical value, 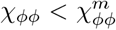 (see SI). In the phase diagrams of case I, the tie-lines connecting coexisting pairs of concentrations generally have negative slope, indicating that the LLPS is segregative, namely, one of the phases is rich in the homotypically-attracting solute and poor in the crowder and the other is poor in the homotypically attracting solute and rich in the crowder.

**Figure 3:**
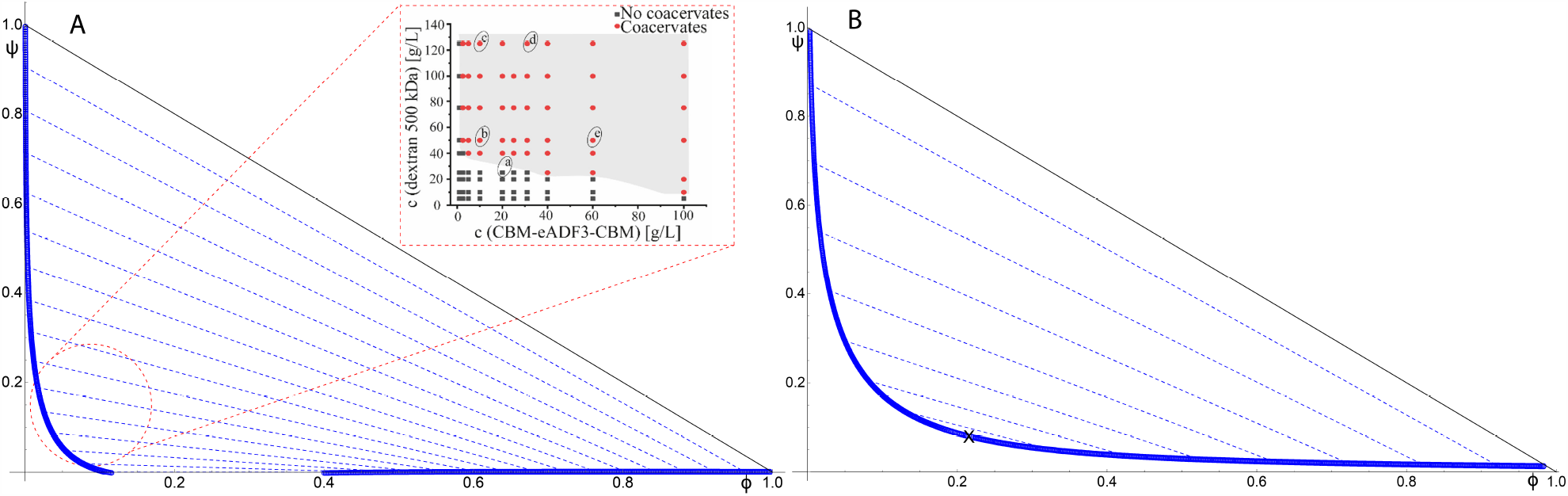
Typical phase diagrams of a ternary system with a homotypically-attracting solute *ϕ* and a crowder *ψ* (where *χ*_*ϕψ*_ = 0 and *χ*_*ψψ*_ = 0). Binodals are marked by blue solid line/open circles, tie-lines by blue, dashed lines, and critical points with “X” marks: (A) Phase diagram for *N*_*ϕ*_ = 10, *N*_*ψ*_ = 6, and *χ*_*ϕϕ*_ = 1.8, which is larger than the critical value 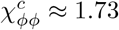 (see SI) required for phase separation in the absence of the crowder. The binodal intersects the line *ψ* = 0, indicating that the solute *ϕ* can phase separate even in the absence of the crowder, *ψ*. However, the presence of crowders (*ψ* > 0) increases the concentration of *ϕ* in the dense phase and decreases it in the dilute phase, rendering LLPS more effective. Inset: in vitro measurements of the experimental phase diagram of such a system, redrawn from Fig. 2 in [31]. The measured phase diagram shows the same trend as the one we predict in the region enclosed in the red, dashed circle. (B) Phase diagram for *N*_*ϕ*_ = 10, *N*_*ψ*_ = 6, and *χ*_*ϕϕ*_ = 1.3 which is less than the critical value 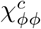 for phase separation in the absence of crowder, but greater than 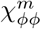 below which the homotypically-attracting solute cannot phase separate even in the presence of crowders (see SI). Within this range of homotypic interactions, the binodal does not intersect the line *ψ* = 0, indicating that crowders are necessary to facilitate LLPS of the homotypically-attracting solute, *ϕ*. In both cases, the presence of crowders cause the tie-lines to be negatively sloped, indicating that the LLPS in the presence of crowders is segregative.

### 4.2 Case II - Heterotypic (*χ*_*ϕϕ*_ = 0, *χ*_*ϕψ*_ > 0): Concentrated phase rich in both solutes (associative LLPS)

For some BMCs, LLPS is the result of heterotypic interactions between molecules of different types, which have only negligible homotypic interactions, so that they do not undergo LLPS when separately mixed with solvent. In the context of our model, we set *χ*_*ϕϕ*_ = *χ*_*ψψ*_ = 0 while keeping *χ*_*ϕψ*_ > 0 to predict the resulting phase diagram (see Fig. 4).

**Figure 4:**
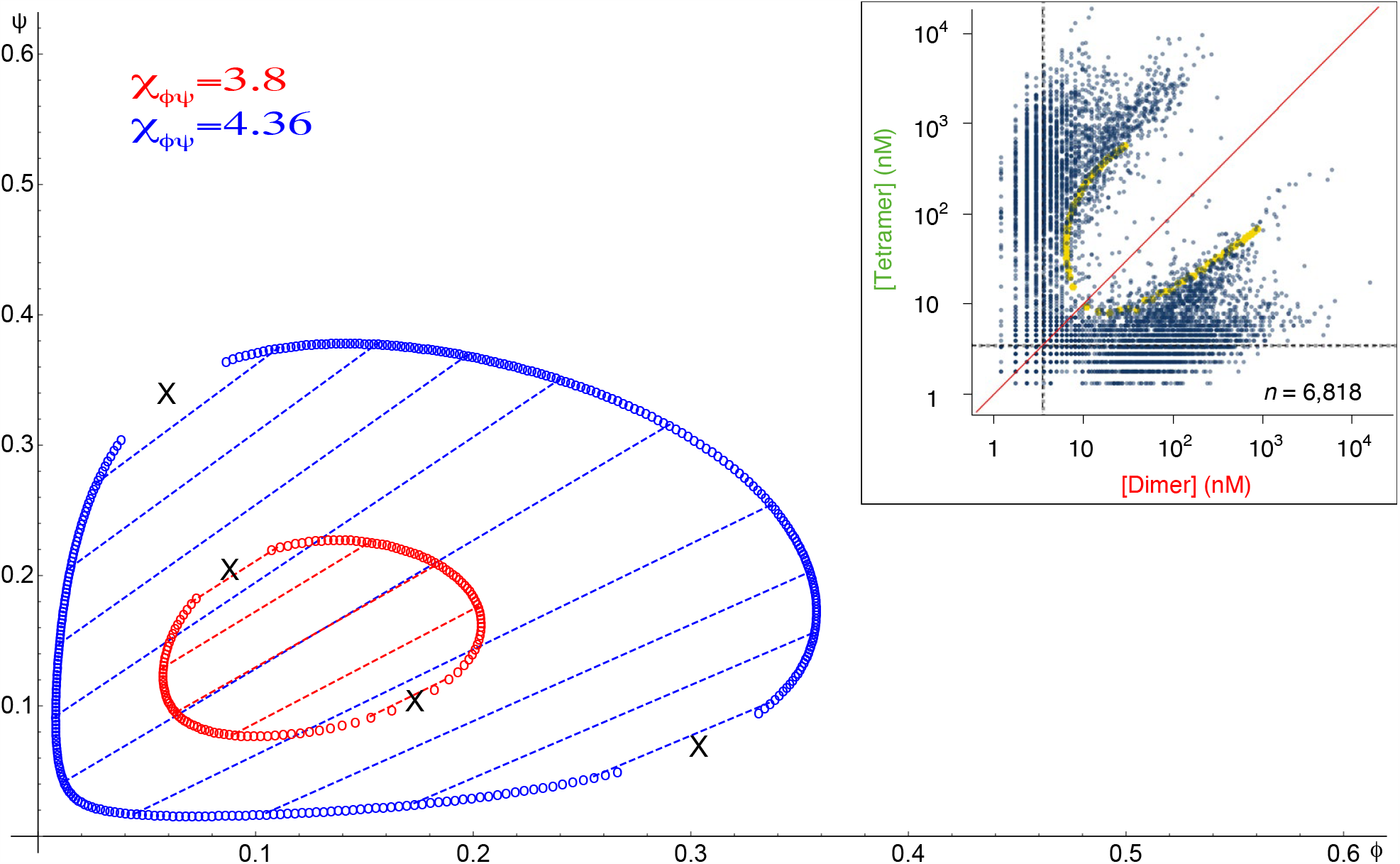
Typical phase diagrams of a ternary system of two solutes that interact purely heterotypically: *χ*_*ϕψ*_ > 0 but *χ*_*ϕϕ*_ and *χ*_*ψψ*_ are both zero. Binodals are marked by solid lines/open circles, tie-lines by dashed lines, and critical points with “X” marks. The red binodal (and tie-lines) is for *χ*_*ϕψ*_ = 3.8 and the blue is for *χ*_*ϕψ*_ = 4.36. In both cases, *N*_*ϕ*_ = 10 and *N*_*ψ*_ = 6. Systems with weaker heterotypic interactions phase separate with smaller binodal loops in the phase diagram as a function of the two concentrations. Both phase diagrams have positively-sloped tie-lines, indicating associative LLPS. Inset: an example of an experimentally measured phase diagram of a system of two multivalent molecules (expressed in living cells) that interact heterotypically. The experimental phase diagram shares the same qualitative features of our predicted phase diagram; figure redrawn from Fig. 2G in [32].

The binodals for different values of the heterotypic interactions, *χ*_*ϕψ*_, form closed loops. The physical interpretation of the loops is that they represent a certain region of “stoichiometric” ratios of the two solutes that interact in a heterotypic manner and strongly enough to phase separate from the aqueous phase. Within these loops, the tie-lines have positive slopes, indicating that the LLPS is associative, namely, one phase is relatively rich in both solutes and the other relative poor in both, but rich in solvent. The area bounded by the loops, which includes the range of protein concentrations that lead to LLPS, shrinks with decreasing *χ*_*ϕψ*_. Below a critical value of heterotypic interaction energy, 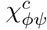, the binodal disappears and the homogenous, mixed state of the solution becomes the equilibrium state for any concentrations *ϕ, ψ*. Our model predicts the value of this critical interaction energy as a function of the solute lengths *N*_*ϕ*_ and *N*_*ψ*_ (see SI).

### 4.3 Case III - Homotypic and heterotypic interactions (*χ*_*ϕϕ*_ > 0, *χ*_*ϕψ*_ > 0): LLPS of a scaffold with a multivalent binding partner

This case accounts for LLPS of multivalent scaffold molecules with homotypic interactions, in the presence of multivalent client (binding partner) molecules that interact heterotypically with the scaffold molecules, but have no homotypic interactions of their own. Within the context of our model of two solutes in a solvent, we set the homotypic interaction of the *ψ* molecules to zero, *χ*_*ψψ*_ = 0, but keep *χ*_*ϕϕ*_, *χ*_*ϕψ*_ > 0; this represents the case where the *ϕ* is the multivalent scaffold and *ψ* is the multivalent client.

The resulting phase diagrams, plotted in Fig. 5, show a surprising, qualitative non-monotonic behavior as the heterotypic interaction parameter is changed. For small heterotypic interactions, the phase diagrams of the scaffold-client model (Fig. 5A) resemble the homotypic phase diagrams of case I (Fig. 3B) with negative-sloped tie-lines. However, as the heterotypic interaction, *χ*_*ϕψ*_, increases, the binodal shrinks and a larger region of the phase diagram is in the single-phase regime (see Fig. 5D), and the slopes of the tie-lines become less negative.

**Figure 5:**
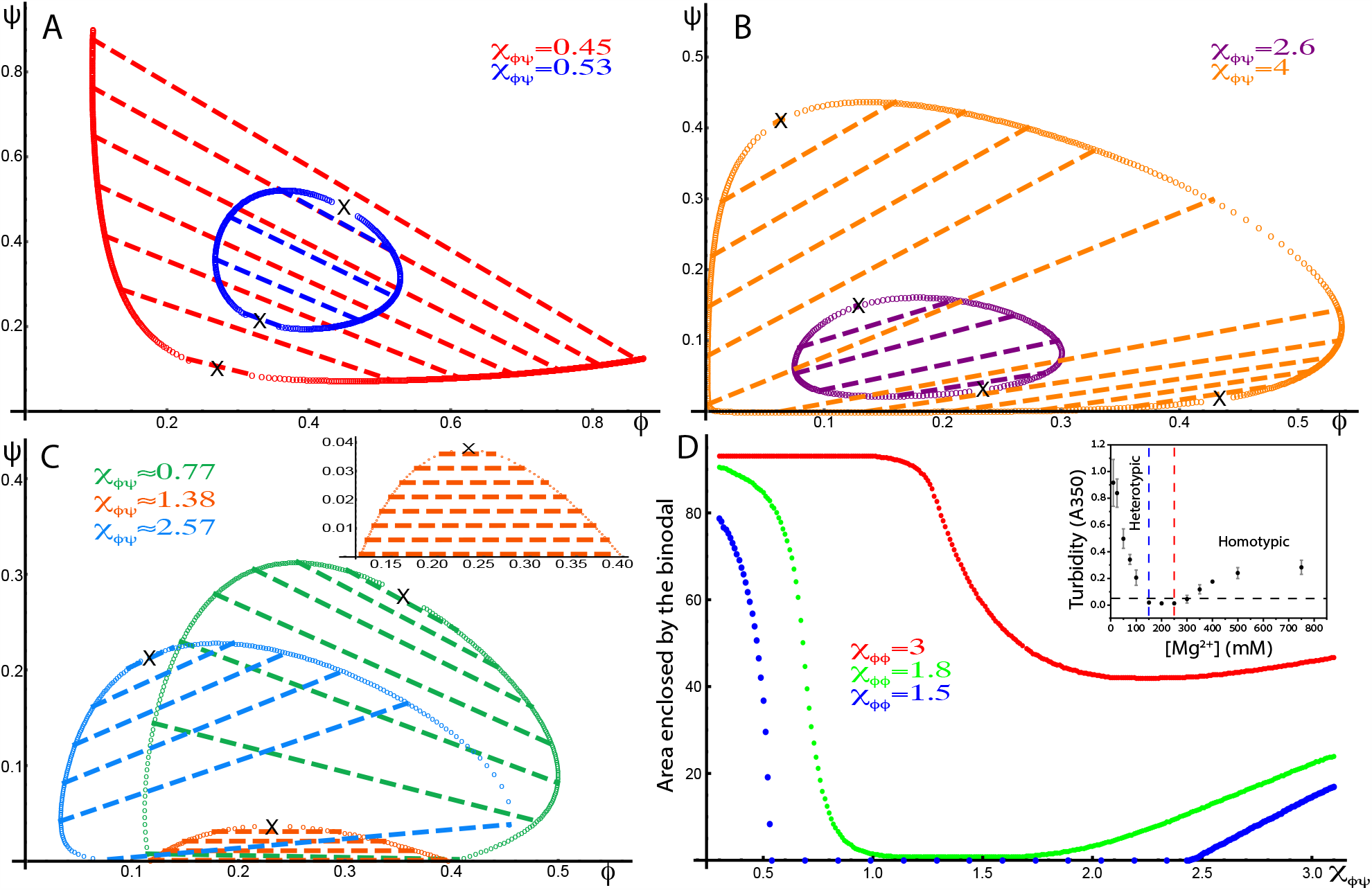
Phase diagrams of ternary systems of a multivalent scaffold (with homotypic interactions) and a multivalent binding partner (that heterotypically associates with the scaffold) in a solvent (*χ*_*ϕϕ*_, *χ*_*ϕψ*_ > 0, *χ*_*ψψ*_ = 0). Binodals are marked by solid lines/open circles, tie-lines by dashed lines, and critical points by “X”. (A+B) Phase diagrams for *N*_*ϕ*_ = 10, *N*_*ψ*_ = 8, and *χ*_*ϕϕ*_ = 1.5 (smaller than 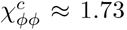), for four different heterotypic interactions, *χ*_*ϕψ*_ = 0.45 (red), *χ*_*ϕψ*_ = 0.53 (blue), *χ*_*ϕψ*_ = 2.6 (purple), and *χ*_*ϕψ*_ = 4 (orange). These diagrams show the discontinuous transition between segregative (negatively-sloped tie-lines) and associative (positively-sloped tie-lines) LLPS as *χ*_*ϕψ*_ increases when 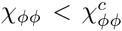. (C) Phase diagrams for *N*_*ϕ*_ = 10, *N*_*ψ*_ = 8, and *χ*_*ϕϕ*_ = 1.8 (larger than 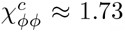), for three different heterotypic interactions, *χ*_*ϕψ*_ = 0.77 (green), *χ*_*ϕψ*_ = 1.38 (orange), and *χ*_*ϕψ*_ = 2.57 (blue). These diagrams show the continuous transition between segregative and associative LLPS as *χ*_*ϕψ*_ increases for the case where there already phase separation in the absence of the binding partner (*ψ* = 1), 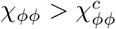. At the critical heterotypic interaction 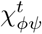 (see SI), the tie-lines are parallel to the *ϕ* axis and the area enclosed by the binodal is minimal (orange and inset). (D) A plot of the area enclosed by the binodal (as percentage of the area enclosed by the triangle in the phase diagram, whose vertex is at the origin and whose base is at the physical limits of the two concentrations, *ϕ* + *ψ* = 1) as a function of *χ*_*ϕψ*_ for three values of *χ*_*ϕϕ*_: *χ*_*ϕϕ*_ = 1.5 (blue), *χ*_*ϕϕ*_ = 1.8 (green), and *χ*_*ϕϕ*_ = 3 (red). When *χ*_*ϕϕ*_ is smaller than 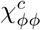 (blue curve), the system exhibits re-entrant behavior indicated by the range of *χ*_*ϕψ*_ for which the area enclosed by the binodal is zero. All three curves show a non-monotonic dependence of the area enclosed by the binodal on *χ*_*ϕψ*_, where the area initially decreases with increasing *χ*_*ϕψ*_, but eventually increases with *χ*_*ϕψ*_. This suggests that the effect of *χ*_*ϕψ*_ on the stability of the LLPS, depends on whether it is associative or segregative. In contrast to the trend as the heterotypic interaction *χ*_*ϕψ*_ is increased, the trend as the homotypic interaction *χ*_*ϕϕ*_ is increased, shows that the area enclosed by the binodal always increases as *χ*_*ϕϕ*_ increases, indicating that homotypic interactions always stabilize LLPS. Inset: Experimental measurements of a phase-separating ternary solution consisting of a positively-charged peptide and RNA in a salt buffer, reproduced from [19]. The turbidity follows the trend predicted for the area of the binodal because the variation of the concentration of M*g*^+2^ ions modulates the balance of homotypic and heterotypic interactions [19], and switches the type of LLPS from associative to segregative, respectively denoted in the inset as “heterotypic” and “homotypic”. of our predicted phase diagram and an experimentally measured one, redrawn from [31] (see Fig. 3A and inset where there is a sharp rise in the binodal near near *ϕ* = 0).

If the scaffold molecule only phase separates in the presence of a crowder (see case I above), the binodal becomes a loop that shrinks with increasing *χ*_*ϕψ*_ and eventually disappears when *χ*_*ϕψ*_ exceeds a lower critical value, 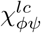, that depends on the homotypic interaction and the molecular sizes (see SI). Further increase of *χ*_*ϕψ*_ above an upper critical value, 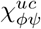, which also depends on the homotypic interaction and the molecular sizes (see SI), results in re-emergence of the binodal as a loop. The re-emerging loop expands in size as *χ*_*ϕψ*_ increases and has positively-sloped tie-lines (see Fig. 5B), similar to case II of the heterotypic phase diagrams (see Fig. 4). For 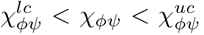, phase separation is completely inhibited and the single-phase state of the solution dominates at all concentrations *ϕ, ψ*, suggesting a re-entrant LLPS in the scaffold-client model as *χ*_*ϕψ*_ increases.

In contrast, if the scaffold has strong homotypic interactions so that it does not require a crowder to undergo LLPS (see Table 2 in the SI), our model predicts a continuous change of the topology of the LLPS with no re-entrant behavior: The area enclosed by the binodal decreases with increasing *χ*_*ϕψ*_ and is minimal at a characteristic value 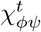 (see SI), and subsequently increases as *χ*_*ϕψ*_ is further increased (see figures 5C and 5D). Similarly, the slope of the tie-lines transition continuously from negative values at 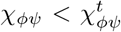 to positive ones for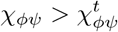 for 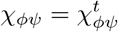 (see Fig. 5C and inset), the tie-lines are completely horizontal.

## 5 Discussion

There are two main differences between a binary solution and one with two or more solutes, which we explain in terms of ternary solutions containing two solutes: First, in the case homotypically-attracting solute in the presence of an inert solute, crowding interactions promote LLPS, and also rotates the tie-lines, which then can make concentration buffering less effective, compared to a binary solution of the homotypically attracting solute and a solvent. Second, the two solutes can undergo LLPS driven by heterotypic interactions as well as homotypic ones. The combination of homotypic and heterotypic interactions can result in continuously varying orientations of the tie-lines. In the subsections below, we discuss the unique features of the phase diagrams of ternary solutions with two solutes that we predicted above, with particular emphasis on the different roles of homotypic vs. heterotypic interactions. We then relate these to their consequence on concentration buffering and review experimental observations that support our theoretical predictions.

### 5.1 Homotypic interactions: Effect of crowders

Our model predicts that addition of a crowder (inert macromolecule) to a homotypically phase-separating solute promotes LLPS. This results in a decrease of both the minimal homotypic interaction strength required for LLPS and the supersaturation concentration of solute for homotypic interaction energies that exceed the minimal one. Furthermore, these effects increase for larger and higher-concentration crowders (see SI for detailed analysis). These predictions are consistent with experimental findings as reflected by the similarities

As shown In Fig. 3, our theory suggests a classification of homotypically phase-separating molecules into two types according to the topology of their phase diagram. The binodal of the first type intersects the line *ψ* = 0 (*ϕ* axis), where *ψ* is the crowder concentration and *ϕ* is the concentration of the phase-separating solute. Phase separation exists even when the crowder concentration is zero. The second type does not intersect this line; the phase separation only exists for finite values of the crowder concentration. This (see SI for further quantification) is equivalent to a known experimental classification of phase-separating molecules into ones that do not require crowding to phase separate (our first type) and ones that do (our second type) [33]; in subsection “Concentration buffering: Noise-ellipse and tie-line orientations” below, we relate this classification to the concentration buffering capabilities of these molecules.

### 5.2 Heterotypic interactions: in the absence or together with homotypic ones

In a phase-separating solution, the different phases can be distinguished as “rich” or “poor” in any given solute by comparing its concentration within that phase to its overall concentration. When the solution has two or more solutes, the LLPS can be classified as “associative” or “segregative” with respect to any pair of solutes if, respectively, each phase is rich and poor in both solutes, or rich in one solute and poor in the other [7]. The phase-diagrams of LLPS that is associative/segregative for the depicted solutes exhibits positively/negatively sloped tie-lines.

Our model predicts that in ternary solutions, LLPS driven by interactions that are either “purely homotypic” (Fig. 3) or “purely heterotypic” (Fig. 4) interactions, is respectively segregative or associative. In addition, we predict an important topological difference between these two cases. For purely homotypic LLPS, the binodal intersects the line describing a state with maximal solute concentration (zero solvent) at the edge of the diagram, characterized by the equation *ϕ* + *ψ* = 1, and possibly the line *ψ* = 0 as well. In contrast, for pure heterotypic LLPS, the binodal takes the form of a closed loop that does not intersect the axes of the phase diagram. This indicates that an excess of either of the two solutes may suppress LLPS or in other words, that LLPS occurs only in a given stoichiometry range. Our generic predictions for the different topologies of the phase diagrams in these two cases, agree with experimental measurements of various specific systems [13, 31, 32, 34]; examples redrawn from [31] and [32] and shown respectively as the insets of figures 3A and 4.

The more intricate scenario for a multi-component solution, in which LLPS is driven by more than one type of interaction, is explored in case III where both homotypic (*χ*_*ϕϕ*_ > 0) and heterotypic (*χ*_*ϕψ*_ > 0) interactions are present. We find that the topologies of the phase diagram and their dependence on the relative strengths of the heterotypic and homotypic interactions can be divided to three classes: (1) 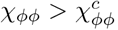, so that *ϕ* can phase-separate even in a binary solution;(2) 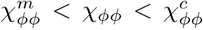, so that *ϕ* can undergo either associative LLPS due to heterotypic interactions or segregative LLPS due to homotypic interactions, but only when enabled by crowding interactions from other solute;(3) 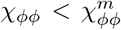, so that the homotypic interactions of *ϕ* are too weak to drive segregative LLPS, and the only LLPS that can occur is associative LLPS driven by heterotypic interactions between *ϕ* and *ψ*. The prediction of re-entrance of the LLPS as the heterotypic interaction is varied in class (2) is supported by in vitro measurements of a ternary solution containing RNA and arginine-rich peptide whose homotypic and heterotypic interactions are continuously modulated by buffer conditions [19] (see Fig. 5D inset, redrawn from [19]). Furthermore, experimental measurements in multiple systems [12, 35, 36, 37, 38, 39, 40], demonstrate transitions between segregative and associative types of LLPS as the interaction strengths are changed by mutations or addition of multivalent binding partners. The latter may may effectively change the interaction strengths [41]. With this interpretation, the experimental measurements of the switch between the associative and segregative types of LLPS are consistent with the predictions of our model for classes (1) and (2).

The areas enclosed by the binodals are plotted in Fig. 5D for classes (1) and (2), with different heterotypic and homotypic interactions, show that an increase of the homotypic interactions promotes LLPS in both types of LLPS. This is because both types of LLPS result in a concentrated phase of the homotypically-attractive solute. In contrast, the effect of changes in the heterotypic interactions generally depends on the type of LLPS. Increase of a heterotypic interaction between two solutes that show associative LLPS promotes the LLPS. Conversely, LLPS that is segregative with respect to two solutes is inhibited by an increase of the heterotypic interaction between the solutes, which tends to associate them. This prediction of potential adverse affect of multivalent binding partners on stability of LLPS is consistent with previous simulations of multicomponent, phase-separating systems [42, 43].

### 5.3 Concentration buffering: Noise-ellipse and tie-line orientations

In section “Concentration buffering in solutions of two solutes”, we explained that LLPS of multi-component solutions can effectively buffer correlated expression noise when the noise-ellipse is oriented in the direction of the tie-lines of the phase diagram. To measure the effects of expression noise, the genes of the proteins of interest can be edited to include two different fused, fluorescent markers. The total, integrated fluorescent signal of each marker within a cell is proportional to the overall concentration of the proteins. Therefore, using calibration curves of the chosen fluorescent markers allows measurements of the overall protein concentrations at different times and of different cells, so that the noise-ellipse can be plotted independent of LLPS.

To discuss the possibility of effective concentration buffering in vivo, we distinguish between BMCs that have relatively high concentrations of only one multivalent macromolecule and those that have relatively high concentrations of more than one multivalent macromolecule. These correspond respectively to the purely homotypic (Case I) and heterotypic (Cases II and III) situations described above for the case of two solutes in a solvent.

In Case I, BMCs are formed by LLPS that is driven by homotypic interactions of one solute and modulated by crowding interactions. In the cellular environment, there are many types molecules that can act as crowders. Since all of these crowders do not interact heterotypically with the solute that form the BMC, they can be treated in an averaged way, as a single, effective crowder. This is supported by many in vitro experiments that recapitulate in vivo crowding using a single type of macromolecule [44]. However, the expression of the effective crowder results from the expression of many, genetically distinct crowders, so that one does not expect any correlation between the expression of the effective, single crowder, and the solute that form the BMC. As a result, the noise-ellipse of the concentration of the solute, *ϕ*, and the effective crowder, *ψ*, is parallel to the *ϕ* axis.

In contrast to the noise-ellipse which are parallel to the *ϕ* axis, our theory predicts that the tie-lines generally are not (see Fig. 3). Only in the specific case where there are no crowders (*ψ* = 0) and the homotypically-attracting solute still phase-separates, are the tie-lines on the *ϕ* axis. However, as the concentration of effective crowder increases, the slope of the tie-lines becomes negative. This means that the effective crowder introduces a mismatch between the direction of the noise-ellipse (parallel to the *ϕ* axis, for the in vivo situation that the effective crowder results from the effects of many, different crowders) and the directions of the tie-lines. Subsequently, the presence of the effective crowder reduces the buffering efficiency in a manner that depends on its concentration. This prediction agrees with the experimental measurements of [16].

In that paper, measurements of the noise in the expression of a homotypically-attracting protein in vivo showed a decrease subsequent to LLPS, but to a lesser degree than predicted for binary solution. A detailed, dynamic theory, predicted trends of the noise reduction as a function of protein concentration, but with higher efficiency than observed (Fig. 2D in [16]). While their theory took into account diffusion of the protein, it did not treat the effect of crowders which, our theory suggests, may account for the reduced buffering efficiency.

If the expression of the genes encoding the different macromolecules that comprise the BMC are regulated by common “master” molecules (e.g. transcription factors), their expression noise may be correlated, which rotates the effective noise-ellipse relative to the *ϕ* axis (see figures 2C and 2D). For the case of the two multivalent macromolecular solutes, a homotypically-attracting solute and a heterotypic binding partner which is not required for the LLPS of the former, our theory predicts that the slope of the tie-lines increases in a continuous manner as the heterotypic interactions are increased. This suggests that for the in vivo situation of many interacting solutes, the orientation of the tie-lines in the phase-diagram changes continuously with the heterotypic interactions of these solutes. In that case, mutations of the various macromolecules that modulate their heterotypic interactions, can change the orientation of the tie-lines in the phase-diagrams. If the effect of the expression noise is deleterious to the organism, then mutations that decrease the angle between the noise-ellipse and the tie-lines may be evolutionary selected, leading to alignment of the orientations of the noise-ellipse and tie-lines over evolutionary timescales, and subsequently to more effective concentration buffering. We note that a recently investigated system provides supporting evidence for this hypothesis: The expressions of two proteins, the homotypically-attracting NPM1 and its multivalent binding partner SURF6, which undergo associative LLPS to form the outer layer of the nucleolus [38], are induced by a common transcription factor, MYC-C [22]. This suggests that the major axis of the noise-ellipse is positively-sloped (see Fig. 2C), and may be aligned with the orientations the tie-lines, which are positively-sloped as well (see blue curve in Fig. 5C). This alignment indicates that concentration buffering is very effective, since the noise mostly changes the concentrations along the tie-lines, resulting in equilibrium concentrations of the two coexisting phases that are always the same, independent of noise except that associated with the minor axis of the ellipse.

In some cases, there are regions of the phase diagrams where the tie-lines are nearly parallel to each other. This means that small variations in the overall concentrations in the direction perpendicular to the tie-lines, result in only small changes to the coexisting, equilibrium concentrations. In other cases, there may be regions where the angles of the tie-lines (and hence the equilibrium concentrations) change more sharply even for relatively small variations of the overall concentrations; this is seen as a “splay” of the tie-lines in the phase diagram. In this case, the splay of the tie-lines serves to amplify the effect of noise on the equilibrium concentrations of the coexisting phase in the side of the phase diagram where the tie-lines are maximally splayed (see Fig. S1 in the SI). This then degrades the concentration buffering. On the other hand, on the opposite side of the phase diagram where the tie-lines converge, the concentrations of the equilibrium phases are even more robustly maintained even in the presence of noise. Thus, on this side of the phase diagram, the effectiveness of concentration buffering may be significantly enhanced due to the proximity of the various tie-lines, even in the presence of a noise-ellipse which is not aligned with the orientation of the tie-lines (see Fig. S1 in the SI). We hypothesize that such large splay may serve - in the case of phases at the side of the phase diagram where the tie-lines converge - as an additional buffering mechanism that protects essential biochemical processes from rare “catastrophic variations” of overall solute concentrations. An example of such a catastrophe is nuclear rupture which may happen during cell migration, resulting in DNA damage and mixing of the cytoplasm and nucleoplasm [45, 46, 47]. Nuclear BMCs such as DNA repair puncta [48], may utilize splay-based buffering to make DNA repair robust to the loss of the diffusion barrier of the nuclear envelope. Future investigations that correlate the phase diagrams of BMCs with such functions is required to explore this hypothesis.

### 5.4 Concluding remarks

In this paper, we formulated theory of two solutes in a solvent to predict the generic behaviors of LLPS in multi-component solutions. We reached the following qualitative conclusions: (1) Crowding effects promote LLPS but decrease its concentration buffering efficiency. (2) The effect on LLPS of heterotypic interactions between two solutes, depends on whether the LLPS is segregative or associative with respect to those solutes; its effect on buffering thus depends on the overall interactions of the entire set of solutes driving the LLPS (interaction network). (3) LLPS involving a subset of the solutes may buffer concentration fluctuations of these solutes if the interaction network of the solutes reflects correlations between the concentration fluctuations. Mutations affecting intermolecular interactions may be subjected to evolutionary selection that improves concentration buffering.

The theory presented here is for two-phase equilibria of two solutes and a solvent. However, we expect the three conclusions of the preceding paragraph to hold for solutions containing many solutes due to the following reasons, as long the LLPS forms two phases, which is true for most biological BMCs: First, since the effect of crowders is independent of any specific chemical characteristic, a solution with many crowder types can be effectively mapped to a solution with a single crowder type, which is treated in our theory. Second, a phase diagram of LLPS of many solutes will be associative or segregative with respect to each pair. This is affected by the overall interaction network. Therefore, it is important to note that the effect on LLPS of a change, e.g. due to a mutation, of heterotypic interaction between a specific pair of solutes depends on the overall interaction network driving the LLPS. Third, as we explain in the SI, in many-solute systems, noise-ellipses and tie-lines can be defined in their multi-dimensional phase diagrams. Since the tie-lines in LLPS of many solutes always define an orientation, even in a multi-dimensional space, as does the major axis of the multidimensional noise-ellipse, our conclusions about concentration buffering when these directions are aligned, are still relevant.

## Acknowledgements

We are grateful to Omar Adame Arana, Gaurav Bajpai, Lucy Brennan, Guoming Gao, Anthony Hyman, Frank Jülicher, Gary Karpen, Adam Klosin, Emmanuel Levy, and Joshua Riback for valuable discussions. The research was supported by the Volkswagen foundation Grant, the Weizmann-Curie Grant, and the Pearlman Grant.

