## Supplementary material for "Physical theory of biological noise buffering by multi-component phase separation"

#### 1 Phase separation of systems with multiple solutes

The equilibrium thermodynamics of a system of $N$ solutes in a solvent, whose concentrations are characterized by the set $\{\phi_1, \cdots, \phi_N\}$, is described by its free energy density function $f(\phi_1, \cdots, \phi_N)$. In equilibrium, the number of coexisting phases and their concentrations are determined by minimization of the total free energy (the spatial integral of $f$) subject to conservation of the number of molecules of each species (including the solvent) and (for an incompressible system), their molecular volumes. If the equilibrium state of the system is a single, mixed phase of all the solutes in the solvent, then the equilibrium concentrations of the solutes are constant (neglecting local and transient thermal fluctuations), and the mean-field total free energy $F$ is related to the free energy density by $F = Vf(\phi_1, \cdots, \phi_N)$, where $V$ is the volume of the entire system. In contrast, if the equilibrium state comprises multiple phases, then the total free energy is the sum of the products of the free energy densities and the volumes of each of the phases. When the interactions between the solutes are short ranged, the concentrations of solutes are constant within the bulk of the phase, and only vary across the interfaces between phases that, except very close to the critical point, have a width on the order of the molecular size. In this case, the total free energy can be approximated as $F = \sum_{j=1}^{k} V_j f(\phi_{1j}, \cdots, \phi_{Nj})$, where $\phi_{ij}$ is the concentration of solute $i$ in phase $j$, $V_j$ is volume of phase $j$ , and $k$ is the number of phases that coexist in equilibrium; the contributions of the interfacial energies to the free energy are not considered in the thermodynamic limit that the system volume is much greater than its cross-sectional area.

The equilibrium state of the system is the one that minimizes the total free energy of the system, subject to the constraints that the total number of each molecular species and (for an incompressible system), the overall volumes of all the molecules (including solvent) in the system are constant. To satisfy these conditions, the free energy is supplemented by terms related to the chemical potentials and osmotic pressure, which results in the grand potential $G$:

$$G = \sum_{j=1}^{k} V_j f(\phi_{1j}, \cdots, \phi_{Nj}) + \sum_{i=1}^{N} \eta_i \left( V\bar{\phi}_i - \sum_{j=1}^{k} V_j \phi_{ij} \right) - \pi \left( V - \sum_{j=1}^{k} V_j \right) \tag{1}$$

where $\bar{\phi}_i$ is the system-averaged, overall concentration of solute $i$, $\eta_i$ is the chemical potential of solute $i$, and $\pi$ is the osmotic pressure of the system. Physically, the chemical potential of a solute represents the change in the free energy due to addition or removal of one molecule of that solute, and the osmotic pressure is the free energy density change due to local changes of volume (expansion or compression). In equilibrium, the chemical potentials and osmotic pressures of the coexisting phases must be equal. Otherwise, the total free energy will not be minimal, since fluxes of solutes or solvent molecules between phases can decrease the total free energy of the system.

Mathematically and equivalently, the equality of chemical potentials $\eta_i$ and osmotic pressure $\pi$ result from the minimization of the grand potential of Eq. 1 with respect to all of its variables: the concentrations of solutes in each of the coexisting phases, $\phi_{ij}$, and the volumes of the different phases, $V_j$. The values of the chemical potentials and osmotic pressure are then determined by the conservation constraints on number of each species and their molecular volumes. This results in the following equations:

$$\frac{\partial f\left(\phi_{1j},\cdots,\phi_{Nj}\right)}{\partial\phi_{ij}}-\eta_i = 0 \quad (2)$$

$$f\left(\phi_{1j},\cdots,\phi_{Nj}\right)-\sum_{i=1}^{N}\eta_i\phi_{ij}+\pi = 0 \quad (3)$$

$$V\bar{\phi}_i-\sum_{j=1}^{k}V_j\phi_{ij} = 0 \quad (4)$$

$$V-\sum_{j=1}^{k}V_j = 0 \quad (5)$$

Eq. 2 implies the equality of the chemical potentials of each solute in each coexisting phase in equilibrium. Eq. 3 implies the equality of the osmotic pressures in each of the coexisting phases in equilibrium. It is important to note that Eqs. 2 and 3 are sufficient to determine the phase diagram of a system, namely the number of phases and coexisting concentrations for different overall concentrations of solutes. However, if one wishes to determine the actual volumes of the coexisting phases, Eqs. 4 and 4 must be used as well. These two equations enforce the conditions that the total molecular volume of each solute and the solvent remain constant (for an incompressible system).

We note that Eqs. 2, 3, 4, and 5, depend on the number of coexisting phases $k$, which is, a-priori, unknown. Furthermore, the single-phase state with $k=1$, $V_1=V$, and $\phi_{i1}=\bar{\phi}_i$, always and trivially solves this set of equations. This implies that a system which shows LLPS can, in general, exhibit several states which solve the non-linear set of Eqs. 2, 3, 4, and 5. Mathematically, this is because this set of equations characterizes the local extrema of the total free energy of the system rather than its global minimum. In principle, to find the global minimum one must obtain all the possible solutions to this set of equations with different values for $k$ that ranges from 1 to $N+1$ (we show in the next section that $k$ cannot exceed $N+1$) and compare their total free energies. This might be a non-trivial computational challenge. However, the convex "hull" surface which represents the minimal, total free energy as a function of the overall concentrations, provides a simple, computational way to find the global minimum for every combination of overall solute concentrations; for detailed explanation of this method, the reader is referred to [1].

In the section below, we predict the nature of the coexisting phases for multicomponent systems at fixed temperature and interactions, by investigating the mathematical structure of these four equations. We compare the number of equations (that originate in the physics) in Eqs. 2 and 3, and the number of unknown concentrations. In some cases, these numbers are not equal which suggests that the concentrations of the coexisting phases in the multicomponent system are not all specified, leading to implicit relations among them for fixed values of the interactions and temperature. In a simple system of one solute that separates into two phases ($N=1$ and $k=2$), the two coexisting concentrations are completely specified at each value of the temperature and interaction strength. However, for the two-solute systems considered in the main text ($N=2$), that phase separate into two phases ($k=2$), the binodals give a continuous range of values of the two solute concentrations that can exist in the two coexisting phases. As we discuss below, there is also a region in the concentration plane where three phases can coexist ($k=3$) for some $N=2$ systems. We show that this equilibrium specifies, for each of the three coexisting phases, only one pair of (or in general, a discrete set of pairs) solute concentrations and one solvent concentration, and not a continuous range as is the case for the region within the binodal of the two phase-equilibria discussed in the main text. The latter arises from what we mean by implicit relations above: In some of the phase separating systems, the physical equations do not specify specific (or a discrete set) of coexisting concentrations, but an entire range. These implicit relations allow us to define the binodal and tie-lines and discuss their geometry for general, $N$-dimensional phase diagrams that characterize the concentrations of $N$ solutes that separate into $k$ phases. We limit our analysis below to the case where the system phase separate, namely $k\geq 2$.

When $k=N+1$ (e.g., three-phase equilibrium for two solutes), we show below that all the concentrations of the coexisting phases are specified for each region in the phase diagram where this equilibrium exists (there may be a discrete number of such regions). In that case, any noise in the concentrations that lies in the region of the $N + 1$ coexisting phases will still give rise to only that single set of solute and solvent concentrations. This represents complete concentration buffering. However, if the noise lies outside of the regions of the $k = N + 1$ coexisting phases, the buffering may not be complete and one must generalize the situation analogous to the alignment of the noise-ellipses and tie-lines for the case of two-solute systems, as discussed in the main text.

### 2 Geometry of phase diagrams of multi-component systems that phase separate into multiple phases

The systems we consider in the main text comprise two solutes and a solvent ($N = 2$), and phase separate into two phases ($k = 2$). However, LLPS in cellular environment is usually more complicated. Biomolecular condensates (BMCs) commonly consist more than two solutes, and many times the proteins involved in LLPS undergo post-translational-modifications, which further increase the number of different species of solutes in the BMC ($N > 2$). Furthermore, some BMCs are formed by LLPS that results in multiple phases ($k > 2$). For example, the nucleolus has a layered structure that consists three distinct layers [2], and its LLPS leads to coexistence within once BMC of three phases (the layers of the nucleolus) that all coexist with a dilute phase of the same proteins ($k = 4$). Therefore, it is important to analyze phase separating systems with $N, k > 2$ that are more complex than ternary systems which phase separate into two phases. In this section, we investigate the geometry of the phase diagrams, binodals, and tie-lines for the general case of multicomponent, phase separating systems, and relate this to concentration buffering in such systems. We begin by examining a few simple examples of systems with two or three solutes and then generalize the discussion using mathematical arguments to specify the geometry of the phase diagrams, binodals, and tie-lines for more complex, multicomponent systems.

#### 2.1 Simple cases of $N = 2, 3$-component system that phase separates into $k = 2, 3$, or 4 coexisting phases

We begin by analyzing the situation of a ternary system ($N = 2$), which phase separates into three ($k = 3$) phases. The situation of $k = 2$ is analyzed quantitatively in the main text for various types of interactions (homotypic and/or heterotypic). At fixed values of the interactions and temperature, the phase diagrams consist of binodal curves (often loops) that enclose a region in the two-dimensional plane of the solute concentrations. However, this is not the case for three-phase coexistence in these systems. In that case, the coexisting phases are specified by three pairs of concentrations $(\phi, \psi)$, which represent six unknown concentrations. These concentrations are related by the six equations for the equality of chemical potentials (Eq. 2), three for $\phi$ and three for $\psi$, and three equations for the equality of osmotic pressures (Eq. 3), a total of nine equations. However, these equations introduce three additional unknowns: two chemical potentials, $\eta_\phi$ and $\eta_\psi$, and an osmotic pressure $\pi$. Therefore, the number of unknowns (six concentrations, two chemical potentials, and one osmotic pressure), is equal to the number of equations (six from Eq. 2 and three from Eq. 3), so that we expect a discrete number of solutions (sometimes only one) for this system of equations. In that case, there are three vertices that define a triangular region of concentration space with three-phase coexistence. Note that each such region determines a single set of solute and solvent concentrations, in contrast to the situation of two-phase equilibrium in these two-solute systems, where there is a continuous line of concentrations that specify the two possible coexisting phases. We call those vertices the branches of a multinodal. In the general case, a branch of the multinodal is the locus of points in the phase diagram that characterizes possible solute concentrations for one coexisting phase; the multinodal of a system that phase separates into $k$ phases has $k$ branches. To further explore how the geometry of the multinodal changes with the number of solutes or phases, we next analyze systems with three solutes. For such systems, the phase phase diagram is three-dimensional since each phase is characterized by three solute concentrations.

In these systems ($N = 3$) that phase separate into two phases ($k = 2$), the number of unknown concentrations is six, three for each of the two phases. Furthermore, there are three unknown chemical potentials, one for each solute type, and one unknown osmotic pressure, as before. These ten unknowns are related by eight equations, two equations for equality of chemical potentials for each of the three solutes, and two equations for the equality of osmotic pressures. Therefore, there are a total of eight equations and ten unknowns. The excess of two unknowns compared to the equations implies that the two branches of the binodal (the number of branches of the multinodal is equal to the number of coexisting phases), have two unspecified concentrations, so that they are two-dimensional surfaces. If the system ($N = 3$) phase separates into three phases ($k = 3$), there are three additional unknowns: the solute concentrations within the additional co-existing phase, and four additional equations, one for each of the three chemical potentials and the single osmotic pressures. Therefore, the excess of unknowns over equations is now one, suggesting that the three branches of the multinodal are one-dimensional curves. Similarly, if the system ($N = 3$) phase separates into four phases ($k = 4$), the number of unknowns and equations becomes equal, so that the four branches of the multinodal consist of zero-dimensional points, similar to the three-phase equilibrium considered in the previous paragraph for ternary systems ($N = 2$). However, now each four-phase region is associated with four concentrations (three solutes and on solvent), and not three as before.

In addition to the geometry of the multinodal, it is important to consider the geometry of the tie-lines that connect coexisting concentrations in the case that the system consists many solutes and phase separates into multiple phases. For such general system, we define "tie-surfaces" as the loci of points in the phase diagram that characterize all the possible combinations of overall (system averaged) solute concentrations, in systems that phase separate into the same set of coexisting phases; as we show below, "tie-surfaces" are lines only in the case of two-phase equilibria ($k = 2$). The possible sets of coexisting phases are described by corresponding points of equal chemical potentials and osmotic pressures on different branches of the multinodal. Once these corresponding points of a specific set are determined, the possible overall solute concentrations in systems that phase separate into this specific set of coexisting phases are determined by varying the volumes in Eq. 4, subject to the constraints of Eq. 5, and positive volumes. If the number of equilibrium phases is two ($k = 2$), than the tie-surfaces connect pairs of corresponding points on the two different branches of the binodal, and are thus lines, which are straight because Eq. 4 in linear in the volumes of the phases. This is true regardless of the number of components $N$. For the case of three-phase equilibria, the tie-surfaces connect three corresponding points and are thus planar triangles, and similarly, for four-phase equilibria, the tie-surfaces are triangular pyramids. As in the case of two-phase equilibria, this is true regardless of the number of components in the system.

In conclusion, we have shown for a few specific examples that an increase of the number of equilibrium phases decreases the dimensionality of the branches of the multinodal but increases their number (which is equal to the number of equilibrium phases). In contrast, increasing the number of solutes increases the dimensionality of the branches of the multinodal, as well as that of the phase diagram itself. Furthermore, we have argued that the geometry of the generalized "tie-surfaces" is determined by the number of equilibrium phases and not by the number of components in the system. In the subsection below, we demonstrate that in the general case of an $N$-component system that phase separates into $k$ phases, the multinodal branches are $(N - k + 1)$-dimensional hyper-surfaces, and the tie-surfaces are $(k - 1)$-dimensional simplexes. Additionally, we briefly discuss how the geometry of the tie-surfaces relates to buffering of expression noise in an $N$-solute system.

#### 2.2 General case of $N$-component system that phase separates into $k$-phases

To mathematically prove the concluding statements of the previous subsection, we count the number of equations and unknowns for a general system of $N$-solutes that phase separates into $k$-phases: Eqs. 2 and 3 respectively provide $kN$ and $k$ different equations (for different values of $i = 1, ..., N$ and $j = 1, .., k$), in terms of a total of $kN + N + 1$ unknowns: $kN$ concentrations, $N$ chemical potentials and 1 osmotic pressure. $N + 1$ of these unknowns, the chemical potentials and osmotic pressure, do not appear in Eqs. 4 and 5. Therefore, it is convenient to eliminate the chemical potentials and osmotic pressure from Eqs. 2 and 3. This is achieved by two mathematical steps. First, the dependence on $\eta_i$ in Eq. 3 for each value of $j$, is eliminated by substituting the expression of $\eta_i$ from Eq. 2 with the same value of $j$ into Eq. 3. Second, we choose an arbitrary value of $j$ that characterize a "reference" phase, and subtract Eqs. 2 and 3 for that value of $j$ from

Eqs. 2 and 3 for all the other values of $j$. It is important to note that the choice of the reference phase is arbitrary and lack any physical significance. For the arbitrary choice of $j = k$ as the reference phase (any other choice is equally valid), this results in the following set of equations:

$$\frac{\partial f\left(\phi_{1j},\cdots,\phi_{Nj}\right)}{\partial \phi_{ij}} = \frac{\partial f\left(\phi_{1k},\cdots,\phi_{Nk}\right)}{\partial \phi_{ik}} \tag{6}$$

$$f\left(\phi_{1j},\cdots,\phi_{Nj}\right) - \sum_{i=1}^{N} \phi_{ij} \frac{\partial f\left(\phi_{1j},\cdots,\phi_{Nj}\right)}{\partial \phi_{ij}} = f\left(\phi_{1k},\cdots,\phi_{Nk}\right) - \sum_{i=1}^{N} \phi_{ik} \frac{\partial f\left(\phi_{1k},\cdots,\phi_{Nk}\right)}{\partial \phi_{ik}} \tag{7}$$

where $i = 1,...,N$, and $j = 1,...,k-1$, because $j = k$ was chosen as the arbitrary value of $j$. This set provides $(k-1)(N+1)$ equations, and has $kN$ unknown concentrations. Thus, there is an excess of $kN - (k-1)(N+1) = N + 1 - k$ unknowns compared with the number of equations.

This general treatment allows us to reach the following conclusions related to the number of equilibrium phases $k$: (1) $k$ cannot exceed the number of components ($N$ solutes + 1 solvent), because then there would be more equations than concentrations so that the system of equations is overdetermined and cannot be solved. This is physically reasonable because $N + 1$ is the maximum number of independent concentrations of the $N+1$ components; additional phases beyond $N+1$ can "be incorporated" within the $N+1$ phases whose free energy densities are the lowest. This would decrease the total free energy and only change the volumes of each of the $N + 1$ coexisting phases of lowest free energy densities but not their solute concentrations. (2) For the special case that the number of phases equals the number of components, $k = N + 1$, the number of equations and unknowns in Eqs. 6 and 7 is equal, so that the coexisting concentrations are determined by these equations alone. This represents the situation where the concentrations in each of the coexisting phases are an inherent property of the interactions of the system, and are independent of the overall (system averaged) concentrations, $\bar{\phi}_i$. In such case the concentration buffering is complete, because systems made with different overall solute concentrations phase separate into phases that always have the same solute concentrations. A single solute in a solvent that can phase separate into two coexisting phases, so that $k = 2$ and $N = 1$, is a system of this kind [3]. (3) If $k < N + 1$, we have more unknowns than equations so that additional equations are needed to uniquely determine the coexisting concentrations. These are available from Eqs. 4 and 5, which respectively include $N$ and 1 equations; these $N + 1$ additional equations then relate the coexisting concentrations to the $k$ different volumes of the $k$ coexisting phases as new unknowns. Therefore, the $N + 1$ Eqs. 4 and 5 supplement the $(k - 1)(N + 1)$ Eqs. 6 and 7 for a total of $k(N + 1)$ equations. This is now equal to the number of unknown concentrations $kN$ in the $k$ coexisting phases, in the set Eqs. 6 and 7, and $k$ unknown coexisting volumes in the set Eqs. 4 and 5, leading to a unique solution of all the coexisting concentrations and all the volumes of the phases.

Although the $(k - 1)(N + 1)$ Eqs. 6 and 7 cannot be solved uniquely since the number of unknown concentrations, $kN$ is larger than the number of equations (when $k - N - 1 < 0$), they nevertheless provide $(k-1)(N+1)$ useful equations that relate the $kN$ concentrations. For each arbitrary choice of a reference phase, the $kN$ concentrations of the entire system is divided to $N$ "reference" concentrations that characterize the reference phase and the remaining $(k-1)N$ concentrations (that we denote as "other concentrations") within the rest of the phases. Since the number of equations provided by Eqs. 6 and 7, $(k-1)(N+1)$, is always larger than the number of the "other" concentrations, $(k-1)N$, a subset of $(k-1)N$ equations out of the total of $(k-1)(N+1)$ equations can be used to implicitly express the "other" concentrations in terms of the reference concentrations.

In that case, these "other" concentrations (as functions of the reference concentrations) can be substituted into the remaining $(k-1)(N+1) - (k-1)N = k - 1$ equations that were not used to relate the "other" and reference concentrations. This yields $k - 1$ implicit expressions that relate the concentrations of any chosen reference phase, and are independent of the "other" concentrations or the overall concentrations. Since any phase can be chosen as the reference phase, a set of $k - 1$ expressions exists for any of the coexisting phases. Even though these implicit expressions usually cannot be written as closed forms, their existence is important since they characterize the different coexisting branches of the multinodal in the phase diagrams of the multi-component system. These $k - 1$ expressions that relate the $N$ solute concentrations within a coexisting phase, when plotted in an $N$-dimensional phase diagram, whose $N$ axes are the $N$ solute concentrations, characterize an $(N-k+1)$-dimensional hyper-surface. The solute concentrations of each of the $k$ phase are characterized by points on the hyper-surface of the phase. Thus, each of the $(N-k+1)$-dimensional hyper-surfaces characterizes a branch of the multinodal. There are a total of $k$ $(N-k+1)$-dimensional hyper-surfaces, one for each coexisting phase; together, they form the high-dimensional multinodal in the general case of a system of a solvent with $N$ solutes that phase separates into $k$ phases.

For any of the points on each of these $(N-k+1)$-dimensional hyper-surfaces, there are $k-1$ corresponding points, one on each of the other surfaces. These sets of $k$ points represent the concentrations of $N$ solutes in $k$ coexisting phases are related by equality of chemical potentials (Eq. 2), and osmotic pressures (Eq. 3). For every set of $k$ coexisting points on the multinodal, it is useful to define the tie-surface as the locus of points in the $N$-dimensional diagram that represents overall solute concentrations of systems that phase separate into phases whose solute concentrations are described by these $k$ points. Once a set of $k$ corresponding points is determined from Eqs. 2 and 3, the tie-surfaces are can be plotted using Eqs. 4 and 5. This is done by varying the volumes of the phases in Eq. 4 under the physical constraint that the volumes are positive and satisfy Eq. 5. Because 4 is linear in the volumes of the phases, then in the general case of $k$ phases, the tie-surfaces are $(k-1)$-simplexes [4].

Furthermore, in the general case that the system has $N$ solutes, the noise-ellipse becomes an $N$-dimensional noise-ellipsoid with $N$ axes. However, since one-dimensional tie-lines characterize phase diagrams of any system that phase separates into two phases, then, even for this general case, there still is a single orientation of the tie-lines that, when aligned with the major axis of the $N$-dimensional noise-ellipsoid, will result in the maximally effective concentration buffering. Therefore, our evolutionary hypothesis presented in the main text, can be simply generalized to this case. When the number of equilibrium phases, $k$, is larger than two, the tie-lines become $(k-1)$-simplexes, and thus have $k-1$ directions in concentration space that buffer concentration fluctuations. This suggests that in addition to the noise associated with the major axis, noise associated with the other $k-2$ additional axes of the noise-ellipsoid can be buffered, and facilitate more effective concentration buffering. Similar to the evolutionary hypothesis presented in the main text for $k=2$, mutations that change the orientations of these $(k-1)$-simplexes in concentration space and result in their alignment with the $k-1$ largest axes of the noise-ellipse, may be evolutionary selected. In the extreme limit that $k=N+1$, the tie-surfaces are $N$-dimensional volumes that in some cases, might fully enclose the noise-ellipsoid, which is also $N$-dimensional. In that situation, the noise may be completely buffered by changes of the volumes of the coexisting phases without any changes of the concentrations of solutes within them. In this case, the coexisting concentrations are independent of the overall concentrations, as indeed was shown above to be the case for $k=N+1$.

### 3 Flory Huggins theory generalized to multiple solutes

So far, we have not specified the free energy density, so that our previous conclusions above are generic to any system of molecules with short-range interactions. However, to determine the phase diagrams for a specific system, an appropriate model for the free energy density must be specified. The focus of this paper is cellular systems. In such systems, the phase separating molecules are usually linear, intrinsically disordered proteins (IDPs) and RNA molecules. Their conformations are similar to those of flexible polymers (and therefore fractal) so that the individual molecules that can interpenetrate [5]. In that case, monomers can interact equally well with each other whether they belong to the same polymer or to different ones. We note that this would not be the case for globular proteins which would only interact via their surface amino-acids and not interpenetrate. This suggests that a mean-field treatment of the interactions of polymers that depend only on the local concentrations of each of the different types of macromolecules in the system is a good approximation for the system. This is precisely the physical picture that underlies the mean-field Flory Huggins (FH) model of polymer mixtures, which uses a lattice model to describe the molecular microstates of the system [6]. The lattice size is typically taken to be of order the persistence length of the chains which we take to be identical. The solvent molecules, which are typically much smaller than the size of one persistence length of the chains, are treated as a "continuum" that fill any volume that is left after placing the two solutes on the lattice. Since, in the lattice model, the smallest volume that can be filled by the "continuum" solvent is one lattice site, the fundamental units of solvent considered in our theory are "blocks" containing many molecules that together occupy one lattice site. In the framework of the FH model, the free energy density for a multicomponent system of $N$ solutes and a solvent is:

$$f = \sum_{i=1}^{N} \frac{\phi_i}{N_i} \log\left(\frac{\phi_i}{N_i}\right) + \phi_s \log(\phi_s) - \frac{1}{2} \sum_{i,j=1}^{N} \tilde{\chi}_{ij}\phi_i\phi_j - \sum_{i=1}^{N} \tilde{\chi}_{is}\phi_i\phi_s - \frac{1}{2}\tilde{\chi}_{ss}\phi_s^2 \tag{8}$$

where $\phi_i$ is the volume fraction of solute $i$, $N_i$ is the size of solute $i$ in units of persistence length, $\phi_s$ is the volume fraction of the solvent, $\tilde{\chi}_{ij}$ is the interaction energy of monomers (persistence length segments of the size of one lattice constant) of solutes $i$ and $j$, $\tilde{\chi}_{is}$ is the interaction energy of a "block" of solvent molecules and a monomer of solute $i$, and $\tilde{\chi}_{ss}$ is the interaction energy of two solvent "blocks" [6]; positive values of $\tilde{\chi}_{ij}$, $\tilde{\chi}_{is}$, and $\tilde{\chi}_{ss}$, represent attractive interactions and negative ones, repulsive interactions. Since the solutes and solvent are the only components in the system, their local volume fractions must sum to unity (within the incompressible lattice model). Therefore, the volume fraction of the solvent can be expressed as $\phi_s = 1 - \sum_{i=1}^{N} \phi_i$. Substitution of this expression into the free energy in Eq. 8, results in the following simplified expression for the FH free energy with the explicit dependence on the solvent volume fraction, $\phi_s$, eliminated:

$$f = \sum_{i=1}^{N} \frac{\phi_i}{N_i} \log(\phi_i) + \left(1 - \sum_{i=1}^{N} \phi_i\right) \log\left(1 - \sum_{i=1}^{N} \phi_i\right) - \frac{1}{2} \sum_{i,j=1}^{N} \chi_{ij}\phi_i\phi_j \tag{9}$$

where constant terms and terms which are linear in the $\phi_i$-s were omitted because they do not affect the LLPS thermodynamics. [Linear terms in the concentration merely renormalize the chemical potentials.]

In Eq. 9, the Flory interaction term $\chi_{ij}$ describes an effective interaction between monomers of solutes $i$ and $j$, relative to their interactions with the solvent, $\chi_{ij} \equiv \tilde{\chi}_{ij} + \chi_{ss} - \chi_{is} - \chi_{js}$. If $i \neq j$, $\chi_{ij}$ describes heterotypic interactions, while if $i = j$, $\chi_{ij}$ describes homotypic interactions. We note that, as captured by the model of stickers and spacers (see section "Theoretical model of multi-component LLPS" in the main text), each phase-separating solute in biological cells can be thought of as a heterogeneous polymer (due to their different amino acid sequences), so that different monomers comprising a given solute are chemically distinct and may in principle, each interact with differing energies. However, in the mean-field spirit of the FH model, the interaction parameters $\chi_{ij}$ are interpreted as interaction energies that were averaged over all the different pairs of monomers of solutes $i$ and $j$.

Substituting the FH free energy into the generic equations for the binodal, Eqs. 2 and 3, results in a set of equations whose solution predicts the concentrations of each species in each of the coexisting phases. These are functions of the solute sizes and interactions:

$$\frac{1}{N_i} \log(\phi_{i\ell}) - \log\left(1 - \sum_{i=1}^{N} \phi_{i\ell}\right) - \sum_{j=1}^{N} \chi_{ij}\phi_{j\ell} + \frac{1}{N_i} - 1 = \eta_i \tag{10}$$

$$\log\left(1 - \sum_{i=1}^{N} \phi_{i\ell}\right) + \sum_{i=1}^{N} \left(1 - \frac{1}{N_i}\right) \phi_{i\ell} + \frac{1}{2} \sum_{i,j=1}^{N} \chi_{ij}\phi_{i\ell}\phi_{j\ell} = -\pi \tag{11}$$

where $i$ and $j$ are indices that denote types of solutes which range from 1 to $N$, and $\ell$ is an index that denotes phases which ranges from 1 to the number of phases $k$. Once the concentrations in each of the coexisting phases are determined from Eqs. 10 and 11, the volumes of the coexisting concentrations are determined by Eqs. 4 and 5 as in the previous sections.

For the case of ternary systems of two solutes in a solvent that is the focus of this paper, Eq. 9 simplifies to yield the following form which we use in the main text:

$$f(\phi, \psi) = \frac{\phi}{N_\phi} \log(\phi) + \frac{\psi}{N_\psi} \log(\psi) + (1 - \phi - \psi) \log(1 - \phi - \psi) - \frac{1}{2}\chi_{\phi\phi}\phi^2 - \chi_{\phi\psi}\phi\psi - \frac{1}{2}\chi_{\psi\psi}\psi^2 \tag{12}$$

where $\phi_1$, $\phi_2$, $N_1$, and $N_2$, are respectively replaced by the simpler notations of $\phi$, $\psi$, $N_\phi$, and $N_\psi$. For the common case of two-phase equilibria, which characterizes most BMCs, the equations for the coexisting concentrations, Eqs. 10 and 11 become:

$$\frac{1}{N_\phi}\log\left(\frac{\phi_1}{\phi_2}\right)-\log\left(\frac{1-\phi_1-\psi_1}{1-\phi_2-\psi_2}\right)-\chi_{\phi\phi}\left(\phi_1-\phi_2\right)-\chi_{\phi\psi}\left(\psi_1-\psi_2\right) = 0 \quad (13)$$

$$\frac{1}{N_\psi}\log\left(\frac{\psi_1}{\psi_2}\right)-\log\left(\frac{1-\phi_1-\psi_1}{1-\phi_2-\psi_2}\right)-\chi_{\phi\psi}\left(\phi_1-\phi_2\right)-\chi_{\psi\psi}\left(\psi_1-\psi_2\right) = 0 \quad (14)$$

$$\log\left(\frac{1-\phi_1-\psi_1}{1-\phi_2-\psi_2}\right)+\left(1-\frac{1}{N_\phi}\right)\left(\phi_1-\phi_2\right)+\left(1-\frac{1}{N_\psi}\right)\left(\psi_1-\psi_2\right) + \quad (15)$$

$$\frac{1}{2}\chi_{\phi\phi}\left(\phi_1^2-\phi_2^2\right)+\chi_{\phi\psi}\left(\phi_1\psi_1-\phi_2\psi_2\right)+\frac{1}{2}\chi_{\psi\psi}\left(\psi_1^2-\psi_2^2\right) = 0 \quad (16)$$

where the subscripts $1, 2$ respectively label the two coexisting phases. This set of equations characterizes the binodal of the system. In order to find the volumes of the coexisting phases for a specific phase-separating system with overall solutes concentrations $\bar{\phi}$ and $\bar{\psi}$, the equations for the conservation of solutes and volume, presented in Eqs. 4 and 5 for the general case, are used to supplement Eqs. 13. In the case of two-phase equilibria, these conservation equations obtain the following form:

$$V_1\phi_1+V_2\phi_2 = V\bar{\phi} \quad (17)$$

$$V_1\psi_1+V_2\psi_2 = V\bar{\psi} \quad (18)$$

$$V_1+V_2 = V \quad (19)$$

### 4 Supplementary results

Motivated by the fact that many BMCs in nature contain no more than a single multivalent scaffold, we limit our discussion to ternary systems where the second solute does not have homotypic interactions, $\chi_{\psi\psi} = 0$, so that only solute $\phi$ may serve as a scaffold. As is apparent from the phase diagrams shown in the main text, ternary systems with $\chi_{\psi\psi} = 0$ can phase separate into two phases, but not three or more. This is consistent with the biological observation that to date, most BMCs are a single, highly concentrated phase that coexists in equilibrium with a dilute phase. A minority of BMCs, such as the complete nucleolus [2], comprise multiple phases, and are outside of the scope of the quantitative analysis presented in this paper. However, as explained in subsection “General case of $N$-component system that phase separates into $k$-phases” above, multi-phase equilibria may be modulated by evolutionary pressures to provide concentration buffering that is even more robust than two-phase equilibria. In this section, we theoretically analyze the different possible scenarios of LLPS in ternary systems with $\chi_{\psi\psi} = 0$, and derive analytical expressions for the critical homotypic and heterotypic interactions mentioned in the main text, that are associated with different topologies of the phase diagrams. Tables 1 and 2 summarize the results of this section. Table 1 lists these critical interactions as functions and the molecular sizes, $N_\phi$ and $N_\psi$, and for some critical heterotypic interactions also of the homotypic interaction $\chi_{\phi\phi}$. Table 2 relates these critical interactions to the various, predicted topologies of the phase-diagrams.

We use two physical insights related to the topologies of the phase diagrams to analytically derive the critical interaction parameters. First, the lines $\phi = 0$, $\psi = 0$, and $\phi + \psi = 1$ that enclose the physically relevant region of the phase diagram, where the volume fractions of the solutes and solvent molecules are all positive, all correspond to different types of binary systems. These are respectively: (i) a system that has no $\phi$ solute, (ii) a system with no $\psi$ solute, (iii) a system that is a blend of $\phi$ and $\psi$ solutes, but with no solvent. Therefore, if the binodal of the ternary system intersects any of these lines, the corresponding binary system undergoes LLPS even without the component whose volume fraction is zero. In that case, the effective interaction energy between the two components of the corresponding binary system must be larger than the critical one predicted by FH theory for binary systems [3].

Second, we note that a system can phase separate if and only if there is a region (termed the spinodal) in the phase diagram where the mixed, uniform state of the system ceases to be a minimum of the free energy density $f$. In that region, a uniform, mixed system becomes unstable and undergoes spinodal decomposition [7]. In other words, the existence of a binodal for LLPS implies the existence of a spinodal region which allows us to use the spinodal as an indicator of LLPS in the following sections.

When phase separation exists, the generalization of the "common tangent" construction [3] that gives rise to binary LLPS in multi-component systems, implies that the free energy surface as function of the concentrations, has a concave (spinodal) region, surrounded, on both sides by a region that is convex. The latter extends to small concentrations where the single-phase must be asymptotically stable due to the dominance of the entropy of mixing at low enough concentrations. The existence of such a spinodal region can also be proven mathematically using a simple *reductio ad absurdum* argument: If the mixed phase is a minimum of the free energy for all possible concentration, then the Hessian matrix $H$ of $f$, defined by

$$H = \begin{pmatrix} \frac{\partial^2 f}{\partial \phi^2} & \frac{\partial^2 f}{\partial \phi \partial \psi} \\ \frac{\partial^2 f}{\partial \phi \partial \psi} & \frac{\partial^2 f}{\partial \psi^2} \end{pmatrix} \tag{20}$$

is positive definite in the entire, multi-dimensional space delineated by the concentration axes that give rise to the phase diagram ("concentration space"). In that case, the free energy density $f$ is everywhere convex. Therefore, the total free energy of a two-phase system is always larger than the total free energy of a one-phase system with the same overall concentrations. This because the average free energy density of the two-phase system is characterized by points along a straight line constructed between two points on the surface described by $f(\phi, \psi)$, which is larger than $f$ itself for the same solute concentrations if $f$ is a convex function. Those two points would represent the coexisting phases of a phase separated system and in the presence of an intervening convex region, binary LLPS cannot be thermodynamically favorable. Conversely, if there is a concave region, the system can undergo spinodal decomposition and thus also phase separate where the two concentrations corresponding to the coexisting phases indeed have lower free energies than any of the single phases along the line connecting them. Therefore, when a phase-separated state of the system, is favorable, there must be a region within the area enclosed by the binodal, in which the single-phase state is unstable and $H$, defined by Eq. 20, has at least one negative eigenvalue. On the boundary of this inner, convex region, namely the spinodal surface, $H$ has at least one vanishing eigenvalue, so that the determinant of $H$ vanishes there.

This correspondence between the binodal and spinodal regions, allows us to use the more easily calculated spinodal as an indicator for the existence of a phase-separated (binodal) regions of the ternary systems. We do this below and analyze each of the three cases considered in the main text separately to derive analytical expressions for the critical values of the heterotypic and homotypic interactions that characterize the different cases and phase diagram topologies.

#### 4.1 Case I - Homotypic ($\chi_{\phi\phi} > 0, \chi_{\phi\psi} = 0$): Concentrated phase rich in one solute (segregative LLPS)

In this case, the interaction parameters of the free energy Eq. 12 obey $\chi_{\phi\phi} > 0$, $\chi_{\phi\psi} = 0$, $\chi_{\psi\psi} = 0$. As explained in the main text, ternary systems in this case comprising solvent, a homotypically attracting-solute and a crowder (which has excluded volume interactions with both the other solute and the solvent but no heterotypic or homotypic interactions), may exhibit three types of phase diagrams, depending on the magnitude of the homotypic interaction: (1) For strong homotypic interaction, $\chi_{\phi\phi} > \chi^c_{\phi\phi}$ (given in Table 1), the homotypically-attracting solute can phase separate from the solvent in the absence of the crowder, so that the binodal in the phase diagram intersects the line $\psi = 0$, which characterizes the absence of crowder. (2) The homotypic interaction is strong enough to facilitate LLPS in the presence of a crowder but too weak to promote LLPS in its absence: $\chi^c_{\phi\phi} > \chi_{\phi\phi}$. However, the system can show LLPS in the presence of a crowder whose maximum concentration obeys $\phi + \psi = 1$, which applies for zero solvent; this constrains $\chi_{\phi\phi} > \chi^m_{\phi\phi}$ (given in Table 1). In that case, the binodal does not intersect the line $\psi = 0$, which characterizes the absence of crowder, but does intersect the line $\phi + \psi = 1$, which characterizes the case where there are only the two solutes and no solvent. (3) The homotypic interaction is too weak to promote LLPS even if the solvent is at its maximal concentration (i.e., solvent is replaced by the crowder), so there is no binodal. This happens for $\chi_{\phi\phi}^{m} > \chi_{\phi\phi}$. Below, we derive analytical expressions for critical interaction parameters, $\chi_{\phi\phi}^{c}$ and $\chi_{\phi\phi}^{m}$, as functions of the sizes of the solute and the crowder, $N_{\phi}$ and $N_{\psi}$.

In order to find $\chi_{\phi\phi}^{c}$, we consider the limiting free energy, $f_{\psi=0}^{I}(\phi)$, of the binary system for the case of zero crowder, $\psi = 0$. Eq. 12 then reduces to:

$$f_{\psi=0}^{I}(\phi) = \frac{\phi}{N_{\phi}} \log(\phi) + (1-\phi)\log(1-\phi) - \frac{1}{2}\chi_{\phi\phi}\phi^{2} \tag{21}$$

Since the critical parameter $\chi_{\phi\phi}^{c}$ distinguishes homotypically-attracting solutes that can undergo LLPS in a binary system from those that cannot, it is equivalent to interaction energy at the critical point of LLPS of the binary system, characterized by the line $\psi = 0$. Therefore, $\chi_{\phi\phi}^{c}$ solves the system of equation that results from equating the second and third derivatives of Eq. 21, with respect to $\phi$, to zero [3]:

$$0 = \frac{1}{N_{\phi}\phi_{c}} + \frac{1}{(1-\phi_{c})} - \chi_{\phi\phi}^{c} \tag{22}$$

$$0 = -\frac{1}{N_{\phi}\phi_{c}^{2}} + \frac{1}{(1-\phi_{c})^{2}} \tag{23}$$

where the value of $\phi_{c}$ that solves these equation is the concentration of the solute $\phi$ at the critical point of the binary system of solute $\phi$ and a solvent. $\chi_{\phi\phi}^{c}$ that solves these equations is given by:

$$\chi_{\phi\phi}^{c} = \frac{\left(\sqrt{N_{\phi}}+1\right)^{2}}{N_{\phi}} \tag{24}$$

Similarly, the homotypic interaction $\chi_{\phi\phi}^{m}$ distinguishes between ternary systems where the binodal intersects the line $\phi+\psi = 1$ and those where the binodal does not. The line $\phi+\psi = 1$ characterizes a binary blend of homotypic solute and client with no solvent. We therefore analyze the free energy density, $f_{\phi+\psi=1}^{I}(\phi)$, of this binary system that is derived from Eq. 12 by substituting $\psi = 1-\phi$:

$$f_{\phi+\psi=1}^{I}(\phi) = \frac{\phi}{N_{\phi}} \log(\phi) + \frac{(1-\phi)}{N_{\psi}} \log(1-\phi) - \frac{1}{2}\chi_{\phi\phi}\phi^{2} \tag{25}$$

As indicated by the intersection of the binodal of the ternary system with the line $\phi + \psi = 1$, a binary blend of $\phi$ and $\psi$ undergoes LLPS only when the homotypic interaction is larger than the value for the critical interaction strength of the blend (with no solvent), given by equating the second and third derivatives of Eq. 21, with respect to $\phi$, to zero. We denote by $\chi_{\phi\phi}^{m}$ this critical value derived from the solution for $\chi_{\phi\phi}$ of the following two equations:

$$0 = \frac{1}{N_{\phi}\phi_{m}} + \frac{1}{N_{\psi}(1-\phi_{m})} - \chi_{\phi\phi}^{m} \tag{26}$$

$$0 = -\frac{1}{N_{\phi}\phi_{m}^{2}} + \frac{1}{N_{\psi}(1-\phi_{m})^{2}} \tag{27}$$

where $\phi_{m}$ is the concentration of solute $\phi$ at the critical point of the binary blend. The solution of this system of equations determines the value of $\chi_{\phi\phi}^{m}$:

$$\chi_{\phi\phi}^{m} = \left(\frac{1}{\sqrt{N_{\phi}}} + \frac{1}{\sqrt{N_{\psi}}}\right)^{2} \tag{28}$$

We note that $\chi_{\phi\phi}^{m}$ decreases with increasing $N_{\psi}$. This indicates that the effect of crowders on LLPS of a homotypically-attracting solute increases with the size of the crowder. Furthermore, $\chi_{\phi\phi}^{m}$ attains two important limiting values when $N_\psi \to \infty$ and $N_\psi = 1$. For $N_\psi \to \infty$, the critical interaction $\chi^m_{\phi\phi}$ approaches $N_\phi^{-1}$. This limiting value of $\chi^m_{\phi\phi}$ is a lower bound on the homotypic interaction of a solute $\phi$ that phase separates in the presence of crowders. This corresponds to an interaction of $k_BT$ (the units in which we measure $\chi_{ij}$) per polymer, which is of size $N_\phi$. Smaller values of the interaction are not effective in generating LLPS since then the entropy per polymer (also of order $k_BT$ per polymer) dominates and promotes mixing in a single, homogeneous phase.

In the other limit, where $N_\psi = 1$, the two critical homotypic interactions, $\chi^c_{\phi\phi}$ and $\chi^m_{\phi\phi}$, which are respectively described by Eqs. 24 and 28, become equal. This special case where the minimal homotypic interaction energy required for solute $\phi$ to phase separate in the presence of crowders, $\chi^m_{\phi\phi}$, is equal to its critical interaction energy required to phase separate in the absence of crowders, $\chi^c_{\phi\phi}$, suggests that the crowders have no effect, at least for $N_\psi = 1$. It is easily seen that this indeed is the case, when we write the free energy for case I, which results from Eq. 12 by substitution of $\chi_{\psi\psi} = \chi_{\phi\psi} = 0$ and $N_\psi = 1$. This can be written in the form:

$$
\begin{aligned}
f(\phi,\psi) &= \frac{\phi}{N_\phi}\log(\phi) + (1-\phi)\log(1-\phi) - \frac{\chi_{\phi\phi}}{2}\phi^2 && (29)\\
&+ (1-\phi)\left(\frac{\psi}{1-\phi}\log\left(\frac{\psi}{1-\phi}\right) + \left(1-\frac{\psi}{1-\phi}\right)\log\left(1-\frac{\psi}{1-\phi}\right)\right) && (30)
\end{aligned}
$$

The first three terms of the right hand side of the equation are the same as the FH free energy of a binary system of solute $\phi$ with an effective solvent of volume fraction $1-\phi$. This effective solvent comprises the aqueous phase and the $\psi$ solute which give rise to a mixing entropy of these two components of the effective solvent given by the last term of Eq. 29. This rewriting is only appropriate for $N_\psi = 1$, but has important physical consequences. Since the driving force for LLPS is the homotypic interaction of $\phi$, only the contribution of the first three terms in Eq. 29 is decreased by LLPS. The contribution of the fourth term in Eq. 29 cannot decrease because that is relevant only to the "internal" properties of the effective solvent. As a matter of fact, if we measure the $\psi$ solvent relative to the volume fraction of the effective solvent $\psi' = \psi/(1-\phi)$, we see that the first three terms of the free energy and independent of the last one (thus can be minimized independently). This demonstrates that indeed, when $N_\psi = 1$, the addition of crowders, which appear only in the fourth term of Eq. 29, has no affect on the phase diagrams of the homotypically-attracting solute $\phi$.

In order to investigate the effect of the crowder size $N_\psi$ for general values of $N_\psi$, we consider the spinodal, which is easier to calculate that the full binodal. The Hessian, $H_I$, of the system is obtained by substitution of the free energy in 12 with $\chi_{\psi\psi} = \chi_{\phi\psi} = 0$, into Eq. 20, is:

$$
H_I = \begin{pmatrix} \frac{1}{N_\phi\phi} + \frac{1}{(1-\phi-\psi)} - \chi_{\phi\phi} & \frac{1}{(1-\phi-\psi)} \\ \frac{1}{(1-\phi-\psi)} & \frac{1}{N_\psi\psi} + \frac{1}{(1-\phi-\psi)} \end{pmatrix}
$$

The spinodal is given by $\det(H_I) = 0$, which results in a quadratic function of $\phi$ whose two roots are expressed as follows:

$$
\phi_\pm = \frac{1}{\chi_{\phi\phi}}\left(\frac{1}{2}\left(\Psi \pm \sqrt{\Psi^2 - \frac{4}{N_\phi}}\right) + \frac{1}{N_\phi}\right) \quad (31)
$$

where $\Psi = \chi_{\phi\phi}(N_\psi - 1)\psi + \chi_{\phi\phi} - 1 - \frac{1}{N_\phi}$ is a parameter related to the concentration of the crowder, its size, and the homotypic interaction strength of the $\phi$ solute and its size, but not to the concentration of $\phi$. These two roots, $\phi_+$ and $\phi_-$, characterize the two branches (large and small concentrations of $\phi$ respectively) of the parabolic spinodal. We note that $\phi_+$ and $\phi_-$ depend on the length of the crowder, $N_\psi$, and its concentration, $\psi$, only implicitly through the parameter $\Psi$. Thus, we conclude that the effects of increasing of the length of the crowder and increasing its concentration are equivalent. Furthermore, our analysis leads to the following conclusions: (1) When $N_\psi = 1$ the expression of $\Psi$ becomes independent of $\psi$, implying that the phase diagram of the homotypically-attracting solute $\phi$ is independent of the concentration of crowder $\psi$, in agreement with our conclusions presented above. (2) $\phi_+$ and $\phi_-$ are respectively monotonically increasing and decreasing function of $\Psi$, so that an increase of the crowder concentration or its length makes the spinodal wider, causing it to cover increasing area of the phase diagram. This indicates that both increasing the concentration of the crowder or its length promote LLPS. (3) In the limit $\Psi \to \infty$, $\phi_- \to (\chi_{\phi\phi} N_\phi)^{-1}$, which suggests that if $\chi_{\phi\phi} N_\phi < 1$, the spinodal is entirely outside of the physical domain of concentration space limited by $\phi, \psi \geq 0$ and $\phi + \psi \leq 1$. This means that the homotypic interactions are too weak to promote LLPS, in agreement with our discussion regarding the limiting value of $\chi^m_{\phi\phi}$.

#### 4.2 Case II - Heterotypic ($\chi_{\phi\phi} = 0, \chi_{\phi\psi} > 0$): Concentrated phase rich in both solutes (associative LLPS)

As described in the main text, the binodals for this case are closed loops in the $\phi$-$\psi$ phase diagram (at constant values of $\chi_{\phi\psi}$), that shrink as the heterotypic interaction strength, $\chi_{\phi\psi}$, decreases. This suggests that there is a critical value of the heterotypic interaction parameter $\chi^c_{\phi\psi}$, below which the binodal disappears; namely, the ternary system does not phase separate. To analytically express $\chi^c_{\phi\psi}$ as a function of the sizes of the two solutes, $N_\phi$ and $N_\psi$, we consider the spinodal of the system, which is easier to calculate. At the heterotypic interaction value of $\chi_{\phi\psi} = \chi^c_{\phi\psi}$, the binodal shrinks to a point, so that, as explained above, the spinodal must shrink to a point as well. Therefore, equating the area of the spinodal to zero provides an equation whose solution is $\chi^c_{\phi\psi}$.

For the heterotypic case, the free energy density is related to 12 by setting $\chi_{\phi\phi} = \chi_{\psi\psi} = 0$. For calculation of the spinodal, we find the Hessian, $H_{II}$, which is obtained by substituting this free energy into Eq. 20:

$$H_{II} = \begin{pmatrix} \frac{1}{N_\phi \phi} + \frac{1}{1-\phi-\psi} & \frac{1}{1-\phi-\psi} - \chi \\ \frac{1}{1-\phi-\psi} - \chi & \frac{1}{N_\psi \psi} + \frac{1}{1-\phi-\psi} \end{pmatrix} \tag{32}$$

The spinodal is the locus of points in the $\phi$-$\psi$ plane defined by $\det(H_{II}) = 0$. In terms of the concentrations of the two solutes, their sizes, and the heterotypic interaction parameter, we obtain the following implicit relation:

$$(1 + \psi(N_\psi - 1)) + ((N_\phi - 1) + N_\psi N_\phi (\chi_{\phi\psi}\psi - (\chi_{\phi\psi} - 2))\chi_{\phi\psi}\psi)\phi + N_\psi N_\phi \chi^2_{\phi\psi}\psi\phi^2 = 0 \tag{33}$$

The left hand size is a quadratic function of $\phi$. The discriminant $\Delta_{II}$, of this quadratic form is:

$$\Delta_{II} = ((N_\phi - 1) + N_\psi N_\phi (\chi_{\phi\psi}\psi - (\chi_{\phi\psi} - 2))\chi_{\phi\psi}\psi)^2 - 4N_\psi N_\phi \chi^2_{\phi\psi}\psi(1 + \psi(N_\psi - 1)) \tag{34}$$

When the concentration of the solute $\psi$ is such that the discriminant is positive, there are two values of $\phi$ that solve Eq. 33 for the given value $\psi$. If the range of values of $\psi$ for which the discriminant is positive, are in the region of physical concentrations $0 \leq \psi \leq 1$, then this range of $\psi$ results in a closed spinodal curve whose two halves are related to the two solutions for $\phi$. The minimal and maximal values of $\psi$ along those curved loops are those where the discriminant goes to zero. This is expected, because the binodal is a closed loop (see Fig. 4 in the main text), so that the spinodal, which is always enclosed by the binodal, must be a closed loop as well. We now determine the endpoint values of $\psi$ (where the discriminant goes to zero), which characterize a transition from positive to negative discriminant. At these endpoints the discriminant vanishes so that they must be roots of Eq. 34. Since Eq. 34 is a quartic polynomial in $\psi$, it has four roots that can be analytically expressed as:

$$\psi_1 = \frac{1}{2}\left(1 - \frac{2}{\chi_{\phi\psi}}\left(1 + \frac{1}{\sqrt{N_\phi}}\right) - \sqrt{\left(1 - \frac{2}{\chi_{\phi\psi}}\left(1 + \frac{1}{\sqrt{N_\phi}}\right)\right)^2 - \frac{4}{N_\psi \chi^2_{\phi\psi}}\left(1 + \frac{1}{\sqrt{N_\phi}}\right)^2}\right) \tag{35}$$

$$\psi_2 = \frac{1}{2}\left(1 - \frac{2}{\chi_{\phi\psi}}\left(1 + \frac{1}{\sqrt{N_\phi}}\right) + \sqrt{\left(1 - \frac{2}{\chi_{\phi\psi}}\left(1 + \frac{1}{\sqrt{N_\phi}}\right)\right)^2 - \frac{4}{N_\psi \chi^2_{\phi\psi}}\left(1 + \frac{1}{\sqrt{N_\phi}}\right)^2}\right) \tag{36}$$

$$\psi_3 = \frac{1}{2}\left(1-\frac{2}{\chi_{\phi\psi}}\left(1-\frac{1}{\sqrt{N_\phi}}\right)-\sqrt{\left(1-\frac{2}{\chi_{\phi\psi}}\left(1-\frac{1}{\sqrt{N_\phi}}\right)\right)^2-\frac{4}{N_\psi\chi_{\phi\psi}^2}\left(1-\frac{1}{\sqrt{N_\phi}}\right)^2}\right) \quad (37)$$

$$\psi_4 = \frac{1}{2}\left(1-\frac{2}{\chi_{\phi\psi}}\left(1-\frac{1}{\sqrt{N_\phi}}\right)+\sqrt{\left(1-\frac{2}{\chi_{\phi\psi}}\left(1-\frac{1}{\sqrt{N_\phi}}\right)\right)^2-\frac{4}{N_\psi\chi_{\phi\psi}^2}\left(1-\frac{1}{\sqrt{N_\phi}}\right)^2}\right) \quad (38)$$

where we used the Solve function of Wolfram Mathematica software to obtain these roots. The values of $\phi$ that correspond to the points along the spinodal at which $\psi$ is minimal or maximal are calculated for these four values of $\psi$ using Eq. 33 are:

$$\phi_1 = \frac{1}{2}\frac{1}{\sqrt{N_\phi}+1}\left(1+\sqrt{\left(1-\frac{2}{\chi_{\phi\psi}}\left(1+\frac{1}{\sqrt{N_\phi}}\right)\right)^2-\frac{4}{N_\psi\chi_{\phi\psi}^2}\left(1+\frac{1}{\sqrt{N_\phi}}\right)^2}\right) \quad (39)$$

$$\phi_2 = \frac{1}{2}\frac{1}{\sqrt{N_\phi}+1}\left(1-\sqrt{\left(1-\frac{2}{\chi_{\phi\psi}}\left(1+\frac{1}{\sqrt{N_\phi}}\right)\right)^2-\frac{4}{N_\psi\chi_{\phi\psi}^2}\left(1+\frac{1}{\sqrt{N_\phi}}\right)^2}\right) \quad (40)$$

$$\phi_3 = -\frac{1}{2}\frac{1}{\sqrt{N_\phi}-1}\left(1+\sqrt{\left(1-\frac{2}{\chi_{\phi\psi}}\left(1-\frac{1}{\sqrt{N_\phi}}\right)\right)^2-\frac{4}{N_\psi\chi_{\phi\psi}^2}\left(1-\frac{1}{\sqrt{N_\phi}}\right)^2}\right) \quad (41)$$

$$\phi_4 = -\frac{1}{2}\frac{1}{\sqrt{N_\phi}-1}\left(1-\sqrt{\left(1-\frac{2}{\chi_{\phi\psi}}\left(1-\frac{1}{\sqrt{N_\phi}}\right)\right)^2-\frac{4}{N_\psi\chi_{\phi\psi}^2}\left(1-\frac{1}{\sqrt{N_\phi}}\right)^2}\right) \quad (42)$$

The first two of these four pairs $(\phi_i, \psi_i)$, represent the points along the spinodal that correspond to the maximal and minimal value of $\psi$, while we find below that the two other pairs represent unphysical values. Since, for $\chi_{\phi\psi} > \chi^c_{\phi\psi}$ the spinodal exists and is contained within the physical region of the concentration space, the first two points satisfy the inequalities $\phi, \psi \geq 0$ and $\phi + \psi \leq 1$ for any value of $\chi_{\phi\psi} > \chi^c_{\phi\psi}$. We examine the last two points by considering their limiting values as $\chi_{\phi\psi} \to \infty$, which shows that $\phi_3$ and $\phi_4$ are negative and therefore are not physically relevant. In conclusion, the points $(\phi_1, \psi_1)$ and $(\phi_2, \psi_2)$ represents the points on the spinodal for which the values of $\psi$ are maximal or minimal. When the heterotypic interaction has the critical value, $\chi_{\phi\psi} = \chi^c_{\phi\psi}$, the spinodal shrinks to a point so that the two points of maximal and minimal $\psi$ become identical. Equating the expressions for these two points, $(\phi_1, \psi_1) = (\phi_2, \psi_2)$ to find where their difference vanishes, yields an equation for the critical value of $\chi_{\phi\psi} = \chi^c_{\phi\psi}$ where the spinodal vanishes, whose solution is:

$$\chi^c_{\phi\psi} = 2\left(1+\frac{1}{\sqrt{N_\psi}}\right)\left(1+\frac{1}{\sqrt{N_\phi}}\right) \quad (43)$$

##### 4.3 Case III - Homotypic and heterotypic interactions ($\chi_{\phi\phi} > 0, \chi_{\phi\psi} > 0$): LLPS of a multivalent scaffold with a multivalent binding partner

As explained in the main text, ternary systems with a combination of both homotypic, $\chi_{\phi\phi}$, and heterotypic interactions, $\chi_{\phi\psi}$, may exhibit either segregative LLPS, driven by homotypic interactions, or associative LLPS, driven by heterotypic interactions. This motivates us to consider three classes of such ternary systems that relate to the transition between segregative to associative LLPS as $\chi_{\phi\psi}$ is increased relative to $\chi_{\phi\phi}$. These three classes differ in the magnitude of their homotypic interactions: (1) Ternary systems in which the homotypic interaction $\chi_{\phi\phi}$ is large enough so that the scaffold $\phi$ can phase separate even in the absence of the multivalent binding partner $\psi$. This indicates that the binodals in the phase diagrams of this class intersect the $\psi = 0$ line for any value of $\chi_{\phi\psi}$. Consequently, this class of systems can phase separate for any value of $\chi_{\phi\psi}$, so that the transition from segregative to associative LLPS is continuous as $\chi_{\phi\psi}$ is increased. (2) Ternary systems in which the homotypic interaction $\chi_{\phi\phi}$ is too small to promote phase separation in the absence of the multivalent binding partner $\psi$, so that those binodals do not intersect the line $\psi = 0$. However, for finite concentrations of $\psi$, the crowding due to the $\psi$ may allow for phase separation. Below, we show that this class of systems may exhibit either segregative or associative LLPS (depending on the values of the heterotypic and homotypic interactions). The transition between the two as $\chi_{\phi\psi}$ is varied involves re-entrant behavior: for small values of $\chi_{\phi\psi}$, LLPS is observed. The region of LLPS then shrinks to zero as $\chi_{\phi\psi}$ in increased (i.e., the system is in a single homogenous phase), but for large enough values, the phase diagram again contains a region of LLPS. This is termed re-entrant behavior of the phase separation. (3) Ternary systems in which the homotypic interaction parameter is too small to promote segregative LLPS even in the presence of the $\psi$. However, these systems may still show associative LLPS when $\chi_{\phi\psi}$ is large enough. Since there is no segragative LLPS in this case, there is no transition between segregative and associative LLPs. In the subsections below, we consider each of these three classes of ternary systems separately, and analytically calculate the critical values of $\chi_{\phi\phi}$ that distinguish these classes, as well as the critical values of $\chi_{\phi\psi}$ that are associated with the transition from segregative to associative LLPS.

##### 4.3.1 Class (1) - Large homotypic interaction: $\phi$ can phase separate in the absence of $\psi$

For this class of systems, the homotypic interactions are large enough so that the scaffold $\phi$ can phase separate in the absence of solute $\psi$. The free energy of the binary system that result from setting $\psi = 0$ is the same as the one described in Eq. 21. Therefore, the critical homotypic interaction above which a ternary system belongs to class (1), is the same as $\chi^c_{\phi\phi}$ defined in Eq. 24. In the limit $\chi_{\phi\psi} \to 0$, systems of case III are reduced to those of case I, and their phase diagrams in this limit must be the similar to the one shown in Fig. 3A in the main text, namely ones in which the binodals intersect both the line $\psi = 0$ and the line $\phi + \psi = 1$. However, the phase diagram shown in Fig. 5C in the main text suggests that, as $\chi_{\phi\psi}$ increases, the area of the binodal shrinks and the binodal ceases to intersect the line $\phi + \psi = 1$. Therefore, there must be a critical heterotypic interaction parameter above which the binodals of class (1) do not intersect the line $\phi + \psi = 1$. Because the line $\phi + \psi = 1$ represents a binary blend of solutes $\psi$ and $\phi$ without solvent, we denote this critical value as $\chi^b_{\phi\psi}$. To calculate the value $\chi^b_{\phi\psi}$, we write the energy of the blend, $f_b(\phi)$, as a function of $\phi$ by substituting $\psi = 1 - \phi$ and $\chi_{\psi\psi} = 0$ into the free energy in Eq. 12, which results in:

$$f_b(\phi) = \frac{\phi}{N_\phi}\log(\phi) + \frac{(1-\phi)}{N_\psi}\log(1-\phi) - \frac{1}{2}\tilde{\chi}\phi^2 \tag{44}$$

where $\tilde{\chi} = \chi_{\phi\phi} - 2\chi_{\phi\psi}$ is the effective (term proportional to $\phi^2$) homotypic interaction energy of the $\phi$ solute in the blend; terms linear in $\phi$ were omitted since they contribute only constant terms to the total free energy that renormalize the chemical potential and do not affect the phase separation properties of the system. The binary blend can phase separate when the binodal of the ternary system intersects the line $\phi + \psi = 1$, and its effective homotypic interaction, $\tilde{\chi}$, is larger than its value at the critical point for LLPS of the binary blend. The solute concentration, $\phi_c$, and effective homotypic interaction, $\tilde{\chi}_c$, at the critical point of the blend are the solutions of $\partial^2 f_b/\partial\phi^2 = 0$ and $\partial^3 f_b/\partial\phi^3 = 0$ [3], which are expressed as follows:

$$0 = \frac{1}{N_\phi\phi_c} + \frac{1}{N_\psi(1-\phi_c)} - \tilde{\chi}_c \tag{45}$$

$$0 = -\frac{1}{N_\phi\phi_c^2} + \frac{1}{N_\psi(1-\phi_c)^2} \tag{46}$$

The critical interaction $\tilde{\chi}_c$ that solves these these equations is:

$$\tilde{\chi}_c = \left(\frac{1}{\sqrt{N_\phi}} + \frac{1}{\sqrt{N_\psi}}\right)^2 \tag{47}$$

Since the effective homotypic interaction energy of the blend is related to the homotypic and heterotypic interactions in the ternary system by $\tilde{\chi} = \chi_{\phi\phi} - 2\chi_{\phi\psi}$, it is larger than $\tilde{\chi}_c$ if $\chi_{\phi\phi} - \left(N_\phi^{-\frac{1}{2}} + N_\psi^{-\frac{1}{2}}\right)^2 > 2\chi_{\phi\psi}$. This indicates that the critical value of the heterotypic interaction energy, $\chi^b_{\phi\psi}$, below which the binodal of a class (1) system intersects the line $\phi + \psi = 1$, is:

$$\chi^b_{\phi\psi} = \frac{1}{2}\chi_{\phi\phi} - \frac{1}{2}\left(\frac{1}{\sqrt{N_\phi}} + \frac{1}{\sqrt{N_\psi}}\right)^2 \tag{48}$$

When the heterotypic interaction $\chi_{\phi\psi}$ is increased beyond $\chi^b_{\phi\psi}$, the LLPS changes continuously from segregative (negatively-sloped tie-lines) to associative (positively-sloped tie-lines). Therefore, a characteristic value of the heterotypic interaction, denoted by $\chi^t_{\phi\psi}$, for which the critical tie-line - i.e., the tangent to the critical point, is parallel to the $\phi$ axis, must exist. In this particular case, the equations that determine the concentrations of solutes $\phi$ and $\psi$ at the critical point [8], are identical with those that determine the critical point in a binary system, where $\partial^2 f/\partial\phi^2 = 0$ and $\partial^3 f/\partial\phi^3 = 0$ [3]. However, in this case the value of $\chi_{\phi\phi}$ is fixed, and the unknowns critical parameters are $\phi_c$ and $\psi_c$. Substitution of the free energy in Eq. 12 with $\chi_{\psi\psi} = 0$ into these two simplified equations results in the following set of equations that characterizes the concentrations of the two solutes at the critical point, $\phi_c$ and $\psi_c$:

$$\begin{aligned}\frac{1}{N_\phi\phi_c} + \frac{1}{(1-\phi_c-\psi_c)} - \chi_{\phi\phi} &= 0 \qquad (49)\\ -\frac{1}{N_\phi\phi_c^2} + \frac{1}{(1-\phi_c-\psi_c)^2} &= 0 \qquad (50)\end{aligned}$$

whose solutions are

$$\begin{aligned}\phi_c &= \frac{\left(1+\frac{1}{\sqrt{N_\phi}}\right)}{\sqrt{N_\phi}\chi_{\phi\phi}} \qquad (51)\\ \psi_c &= 1 - \frac{\left(1+\frac{1}{\sqrt{N_\phi}}\right)^2}{\chi_{\phi\phi}} \qquad (52)\end{aligned}$$

These values of the concentrations at the critical point are correct only when the heterotypic interaction attains a the characteristic value, $\chi_{\phi\psi} = \chi^t_{\phi\psi}$, which we now calculate. To relate the heterotypic interaction $\chi^t_{\phi\psi}$ to the concentrations of the solutes at the critical point, we use the fact that the critical point must lie on the spinodal line [8]. Therefore, for $\phi = \phi_c$, $\psi = \psi_c$ (where $\phi_c$ and $\psi_c$ respectively given by Eqs. 51 and 52), and $\chi_{\phi\psi} = \chi^t_{\phi\psi}$, the determinant of the Hessian $H_{III}$ in this case, must vanish. For a ternary system of case III, where homotypic and heterotypic interactions are both present, the free energy is given by substitution of $\chi_{\psi\psi} = 0$ in Eq. 12. The Hessian is then obtained by substitution of this free energy into Eq. 20, which results in the following expression:

$$H_{III} = \begin{pmatrix} \frac{1}{N_\phi\phi} + \frac{1}{1-\phi-\psi} - \chi_{\phi\phi} & \frac{1}{1-\phi-\psi} - \chi_{\phi\psi} \\ \frac{1}{1-\phi-\psi} - \chi_{\phi\psi} & \frac{1}{N_\psi\psi} + \frac{1}{1-\phi-\psi} \end{pmatrix} \tag{53}$$

When $\chi_{\phi\psi} = \chi^t_{\phi\psi}$, and the expressions from Eqs. 51 and 52 are used for $\phi$ and $\psi$, setting $\det(H_{III}) = 0$ results in an equation, whose solution gives the explicit expression of $\chi^t_{\phi\psi}$ as a function of the solute sizes and the homotypic interaction:

$$\chi^t_{\phi\psi} = \frac{\chi_{\phi\phi}}{\left(1+\frac{1}{\sqrt{N_\phi}}\right)} \tag{54}$$

For $\chi_{\phi\psi} < \chi^t_{\phi\psi}$, the tie-lines are negatively-sloped (segregative LLPS), while for $\chi_{\phi\psi} > \chi^t_{\phi\psi}$, the tie-lines are positively-sloped (associative LLPS). Because an increase of the heterotypic interaction between $\phi$ and $\psi$ favor LLPS in which the concentrated phase has considerable amounts of both solutes, the heterotypic interactions destabilize segregative LLPS and stabilize associative LLPS, as indicated by their effect on the area of the binodals (see Fig. 5D in the main text). Therefore, the area of the binodal when the heterotypic interaction parameter is equal to the characteristic one, $\chi_{\phi\psi} = \chi^t_{\phi\psi}$, is expected to be minimal.

###### 4.3.2 Class (2) - Intermediate homotypic interaction: $\phi$ can phase separate segregatively but requires crowding interactions

For ternary systems of this class, the homotypic interaction $\chi_{\phi\phi}$ is too small to promote LLPS in binary systems of the solute $\phi$ and a solvent, but is large enough to drive LLPS when assisted by the crowding interactions with finite concentrations of the multivalent binding partner $\psi$. Consequently, the binodals of systems in class (2) do not intersect the line $\psi = 0$. Therefore, in this case, $\chi_{\phi\phi}$ is smaller than the critical homotypic interaction $\chi^c_{\phi\phi}$ given by Eq. 24, but must be larger than a lower bound, below which, $\phi$ cannot phase separate homotypically at all. Since, as explained at the end of the previous subsection, heterotypic interactions of $\phi$ with $\psi$ inhibit segregative LLPS, $\psi$ is most effective in promoting segregative LLPS when $\chi_{\phi\psi} \to 0$, namely when $\psi$ is a crowder (with only excluded volume interactions with $\phi$ and the solvent) rather than a multivalent binding partner (with non-zero attraction to $\phi$ for $\chi_{\phi\psi} > 0$). Therefore, the lower bound for the homotypic interactions below which the solute $\phi$ cannot undergo segregative LLPS at all, even when promoted by crowding interactions of $\psi$, is the critical value $\chi^m_{\phi\phi}$ derived for case I in Eq. 28; $\chi^m_{\phi\phi}$ is the minimal (critical) homotypic interaction required for phase separation of the binary blend of $\phi$ and $\psi$ in the absence of solvent, i.e., along the line $\phi + \psi = 1$. In conclusion, the homotypic interactions of class (2) systems must lie within the range $\chi^c_{\phi\phi} > \chi_{\phi\phi} > \chi^m_{\phi\phi}$.

We note that this interaction range $\chi^c_{\phi\phi} > \chi_{\phi\phi} > \chi^m_{\phi\phi}$ corresponds to type (2) phase diagrams of systems considered in case I (where $\chi_{\phi\psi} = 0$), whose binodals intersect the line $\phi + \psi = 1$ but not the line $\psi = 0$ (Fig. 3B in the main text). Therefore, in the limiting case $\chi_{\phi\psi} \to 0$, the phase diagrams of class (2) systems must coincide with type (2) diagrams of case I. However, because heterotypic interactions destabilize segregative LLPS, an increase of $\chi_{\phi\psi}$ from zero causes the binodal to shrink, and to eventually become a closed curve that does not intersect the line $\phi + \psi = 1$ (see Fig. 5A in the main text). This is similar to the behavior of class (1) phase-diagrams, whose binodals no longer intersect the line $\phi + \psi = 1$ when $\chi_{\phi\psi}$ exceeds the critical value $\chi^b_{\phi\psi}$. Moreover, the derivation of $\chi^b_{\phi\psi}$ for class (1) systems above, does not require the value of $\chi_{\phi\phi}$ to be within the range that characterize LLPS for class (1) systems. Therefore, the critical value $\chi^b_{\phi\psi}$ that given by Eq. 48, is also the critical heterotypic interaction below (above) which binodals of class (2) systems intersect (do not intersect) the line $\phi + \psi = 1$. Surprisingly, in contrast to class (1) phase-diagrams, a further increase of $\chi_{\phi\psi}$ in class (2) phase diagrams causes the system to exhibit re-entrant behavior. This is because the binodal loop of class (2) systems shrinks with increasing values of $\chi_{\phi\psi}$ and eventually disappear when $\chi_{\phi\psi}$ reaches a critical value $\chi^{lc}_{\phi\psi}$ (see below for its derivation). For larger values of $\chi_{\phi\psi} > \chi^{uc}_{\phi\psi}$ (see Fig. 5B in the main text and below for its derivation), the binodal loop reappears. The phase separation is now associative instead of segregative and the loop expands as $\chi_{\phi\psi}$ is increased beyond the value of $\chi^{uc}_{\phi\psi}$. When the heterotypic interaction is in between these two critical values, $\chi^{lc}_{\phi\psi} < \chi_{\phi\psi} < \chi^{uc}_{\phi\psi}$, LLPS does not occur and the mixed phase of the ternary system is the equilibrium state for any concentrations of the solutes $\phi$ and $\psi$.

To calculate these two critical values, $\chi^{lc}_{\phi\psi}$ and $\chi^{uc}_{\phi\psi}$, as functions of the homotypic interaction $\chi_{\phi\phi}$ and the solute sizes, $N_\phi$ and $N_\psi$, we determine the values of the heterotypic interaction strength at which the spinodal vanishes and then reappears. As explained at the beginning of the "Supplementary results" section above, the binodal region must enclose a spinodal region, so that the values of $\chi_{\phi\psi}$ for which the spinodal vanishes and reappears must be the same as $\chi^{lc}_{\phi\psi}$ and $\chi^{uc}_{\phi\psi}$, respectively. Furthermore, since the binodal curves are closed loops prior to their vanishing and after their reappearance, the spinodal must also be a closed loop enclosed in the binodal region.

The spinodal is defined by the condition that $\det(H_{III}) = 0$ (where $H_{III}$ is given by Eq. 53), which is expressed in terms of the solutes concentrations, their sizes, and the two interaction parameters, as follows:

$$\begin{aligned} 0 &= N_\phi \left(\chi_{\phi\phi} + \chi_{\phi\psi}^2 N_\psi \psi\right)\phi^2 &(55)\\ &+ \left((N_\phi - 1) - N_\phi \chi_{\phi\phi}\left(1 + (N_\psi - 1)\psi\right) + N_\phi N_\psi \chi_{\phi\psi}\psi\left(2 - \chi_{\phi\psi}(1-\psi)\right)\right)\phi + \left(1 + (N_\psi - 1)\psi\right) &(56)\end{aligned}$$

We now analyze the spinodal, in a manner similar to that used in subsection "Case II - Heterotypic ($\chi_{\phi\phi} = 0, \chi_{\phi\psi} > 0$): Concentrated phase rich in both solutes (associative LLPS)". This spinodal is a quadratic function of $\phi$, whose discriminant, $\Delta_{III}(\psi)$, is:

$$\begin{aligned} \Delta_{III}(\psi) &= \left((N_\phi - 1) - N_\phi \chi_{\phi\phi}\left(1 + (N_\psi - 1)\psi\right) + N_\phi N_\psi \chi_{\phi\psi}\psi\left(2 - \chi_{\phi\psi}(1-\psi)\right)\right)^2 &(57)\\ &- 4N_\phi\left(1 + (N_\psi - 1)\psi\right)\left(\chi_{\phi\phi} + \chi_{\phi\psi}^2 N_\psi \psi\right) &(58)\end{aligned}$$

There are two real concentrations $\phi$ that solve the quadratic equation 55, as long as the concentration $\psi$ that appear in Eq. 55 gives a positive discriminant: $\Delta_{III}(\psi) > 0$. This means that for any value of $\psi$ where $\Delta_{III}(\psi) > 0$, there are two points (due to the two solutions of the quadratic equation for the spinodal) on the spinodal that are associated with this value of $\psi$; those two points generally have different values of $\phi$. Thus, continuous regions of $\psi$ that satisfy the inequality $\Delta_{III}(\psi) \geq 0$, form closed curves in the $\phi$-$\psi$ plane, whose maximal and minimal values of $\psi$ are those that satisfy the equality $\Delta_{III}(\psi) = 0$. If these regions, and their corresponding values of $\phi$, (calculated from 55), are within the "physical" range of the two concentrations, $\phi, \psi \geq 0$ and $\phi + \psi \leq 1$, they determine closed, spinodal curves. Below, we determine the roots of the equation $\Delta_{III}(\psi) = 0$, and calculate their corresponding values of $\phi$ using Eq. 55. We then select the "physical" roots that correspond to the the maximal and minimal values of $\psi$ along the spinodal, whose corresponding $(\phi, \psi)$ points are in the region $\phi, \psi \geq 0$ and $\phi + \psi \leq 1$. Since Eq. 57 is a quartic polynomial in $\psi$, it has four roots that can be analytically expressed as:

$$\begin{aligned} \psi_1 &= \frac{1}{2}\left(1 + \frac{\chi_{\phi\phi}}{\chi_{\phi\psi}^2}\left(1 - \frac{1}{N_\psi}\right) - \frac{2}{\chi_{\phi\psi}}\left(1 + \frac{1}{\sqrt{N_\phi}}\right)\right) &(59)\\ &- \frac{1}{2}\sqrt{\left(1 + \frac{\chi_{\phi\phi}}{\chi_{\phi\psi}^2}\left(1 - \frac{1}{N_\psi}\right) - \frac{2}{\chi_{\phi\psi}}\left(1 + \frac{1}{\sqrt{N_\phi}}\right)\right)^2 - \frac{4}{\chi_{\phi\psi}^2}\frac{1}{N_\psi}\left(\left(1 + \frac{1}{\sqrt{N_\phi}}\right)^2 - \chi_{\phi\phi}\right)} &(60)\\ \psi_2 &= \frac{1}{2}\left(1 + \frac{\chi_{\phi\phi}}{\chi_{\phi\psi}^2}\left(1 - \frac{1}{N_\psi}\right) - \frac{2}{\chi_{\phi\psi}}\left(1 + \frac{1}{\sqrt{N_\phi}}\right)\right) &(61)\\ &+ \frac{1}{2}\sqrt{\left(1 + \frac{\chi_{\phi\phi}}{\chi_{\phi\psi}^2}\left(1 - \frac{1}{N_\psi}\right) - \frac{2}{\chi_{\phi\psi}}\left(1 + \frac{1}{\sqrt{N_\phi}}\right)\right)^2 - \frac{4}{\chi_{\phi\psi}^2}\frac{1}{N_\psi}\left(\left(1 + \frac{1}{\sqrt{N_\phi}}\right)^2 - \chi_{\phi\phi}\right)} &(62)\\ \psi_3 &= \frac{1}{2}\left(1 + \frac{\chi_{\phi\phi}}{\chi_{\phi\psi}^2}\left(1 - \frac{1}{N_\psi}\right) - \frac{2}{\chi_{\phi\psi}}\left(1 - \frac{1}{\sqrt{N_\phi}}\right)\right) &(63)\\ &- \frac{1}{2}\sqrt{\left(1 + \frac{\chi_{\phi\phi}}{\chi_{\phi\psi}^2}\left(1 - \frac{1}{N_\psi}\right) - \frac{2}{\chi_{\phi\psi}}\left(1 - \frac{1}{\sqrt{N_\phi}}\right)\right)^2 - \frac{4}{\chi_{\phi\psi}^2}\frac{1}{N_\psi}\left(\left(1 - \frac{1}{\sqrt{N_\phi}}\right)^2 - \chi_{\phi\phi}\right)} &(64)\\ \psi_4 &= \frac{1}{2}\left(1 + \frac{\chi_{\phi\phi}}{\chi_{\phi\psi}^2}\left(1 - \frac{1}{N_\psi}\right) - \frac{2}{\chi_{\phi\psi}}\left(1 - \frac{1}{\sqrt{N_\phi}}\right)\right) &(65)\\ &+ \frac{1}{2}\sqrt{\left(1 + \frac{\chi_{\phi\phi}}{\chi_{\phi\psi}^2}\left(1 - \frac{1}{N_\psi}\right) - \frac{2}{\chi_{\phi\psi}}\left(1 - \frac{1}{\sqrt{N_\phi}}\right)\right)^2 - \frac{4}{\chi_{\phi\psi}^2}\frac{1}{N_\psi}\left(\left(1 - \frac{1}{\sqrt{N_\phi}}\right)^2 - \chi_{\phi\phi}\right)} \Rightarrow &(66)\end{aligned}$$

where we used the Solve function of Wolfram Mathematica software to obtain these roots. The corresponding values of $\phi$ that are calculated for these four values of $\psi$ using Eq. 55 are:

$$\phi_1 = \frac{1}{2}\frac{1}{\sqrt{N_\phi}+1-\frac{\chi_{\phi\phi}}{\chi_{\phi\psi}}\sqrt{N_\phi}}\left(1-\frac{\chi_{\phi\phi}}{\chi_{\phi\psi}^2}\left(1-\frac{1}{N_\psi}\right)\right) \tag{67}$$

$$+ \frac{1}{2}\frac{\sqrt{\left(1+\frac{\chi_{\phi\phi}}{\chi_{\phi\psi}^2}\left(1-\frac{1}{N_\psi}\right)-\frac{2}{\chi_{\phi\psi}}\left(1+\frac{1}{\sqrt{N_\phi}}\right)\right)^2-\frac{4}{\chi_{\phi\psi}^2}\frac{1}{N_\psi}\left(\left(1+\frac{1}{\sqrt{N_\phi}}\right)^2-\chi_{\phi\phi}\right)}}{\sqrt{N_\phi}+1-\frac{\chi_{\phi\phi}}{\chi_{\phi\psi}}\sqrt{N_\phi}} \tag{68}$$

$$\phi_2 = \frac{1}{2}\frac{1}{\sqrt{N_\phi}+1-\frac{\chi_{\phi\phi}}{\chi_{\phi\psi}}\sqrt{N_\phi}}\left(1-\frac{\chi_{\phi\phi}}{\chi_{\phi\psi}^2}\left(1-\frac{1}{N_\psi}\right)\right) \tag{69}$$

$$- \frac{1}{2}\frac{\sqrt{\left(1+\frac{\chi_{\phi\phi}}{\chi_{\phi\psi}^2}\left(1-\frac{1}{N_\psi}\right)-\frac{2}{\chi_{\phi\psi}}\left(1+\frac{1}{\sqrt{N_\phi}}\right)\right)^2-\frac{4}{\chi_{\phi\psi}^2}\frac{1}{N_\psi}\left(\left(1+\frac{1}{\sqrt{N_\phi}}\right)^2-\chi_{\phi\phi}\right)}}{\sqrt{N_\phi}+1-\frac{\chi_{\phi\phi}}{\chi_{\phi\psi}}\sqrt{N_\phi}} \tag{70}$$

$$\phi_3 = -\frac{1}{2}\frac{1}{\sqrt{N_\phi}-1-\frac{\chi_{\phi\phi}}{\chi_{\phi\psi}}\sqrt{N_\phi}}\left(1-\frac{\chi_{\phi\phi}}{\chi_{\phi\psi}^2}\left(1-\frac{1}{N_\psi}\right)\right) \tag{71}$$

$$- \frac{1}{2}\frac{\sqrt{\left(1+\frac{\chi_{\phi\phi}}{\chi_{\phi\psi}^2}\left(1-\frac{1}{N_\psi}\right)-\frac{2}{\chi_{\phi\psi}}\left(1-\frac{1}{\sqrt{N_\phi}}\right)\right)^2-\frac{4}{\chi_{\phi\psi}^2}\frac{1}{N_\psi}\left(\left(1-\frac{1}{\sqrt{N_\phi}}\right)^2-\chi_{\phi\phi}\right)}}{\sqrt{N_\phi}-1-\frac{\chi_{\phi\phi}}{\chi_{\phi\psi}}\sqrt{N_\phi}} \tag{72}$$

$$\phi_4 = -\frac{1}{2}\frac{1}{\sqrt{N_\phi}-1-\frac{\chi_{\phi\phi}}{\chi_{\phi\psi}}\sqrt{N_\phi}}\left(1-\frac{\chi_{\phi\phi}}{\chi_{\phi\psi}^2}\left(1-\frac{1}{N_\psi}\right)\right) \tag{73}$$

$$+ \frac{1}{2}\frac{\sqrt{\left(1+\frac{\chi_{\phi\phi}}{\chi_{\phi\psi}^2}\left(1-\frac{1}{N_\psi}\right)-\frac{2}{\chi_{\phi\psi}}\left(1-\frac{1}{\sqrt{N_\phi}}\right)\right)^2-\frac{4}{\chi_{\phi\psi}^2}\frac{1}{N_\psi}\left(\left(1-\frac{1}{\sqrt{N_\phi}}\right)^2-\chi_{\phi\phi}\right)}}{\sqrt{N_\phi}-1-\frac{\chi_{\phi\phi}}{\chi_{\phi\psi}}\sqrt{N_\phi}} \tag{74}$$

It is important to note that, because of the discontinuous, re-entrant behavior of the phase-diagram, the two “physical” points $(\phi, \psi)$ of the four pairs of solutions are not necessarily the same for the separate continuous domains $\chi_{\phi\psi} > \chi_{\phi\psi}^{uc}$ and $\chi_{\phi\psi} < \chi_{\phi\psi}^{lc}$. We thus consider each domain separately and select the two points that are physical.

For values of $\chi_{\phi\psi} > \chi_{\phi\psi}^{uc}$, the binodal and spinodal form closed loops, whose maximal and minimal values of $\psi$ and their corresponding values of $\phi$, are all in the physical region of the concentration space for any value of $\chi_{\phi\psi} > \chi_{\phi\psi}^{uc}$ (see Fig. 5B in the main text). Therefore, we consider the asymptotic limit, $\chi_{\phi\psi} \to \infty$ for which the expressions for the various sets of values of $\psi$ and $\phi$ detailed above are greatly simplified, and use these simplified expressions to select the values that are physical. In the limit $\chi_{\phi\psi} \to \infty$, $(\phi_1, \psi_1) \to \left(\left(\sqrt{N_\phi}+1\right)^{-1}, 0^+\right)$, $(\phi_2, \psi_2) \to (0^+, 1^-)$, $(\phi_3, \psi_3) \to \left(-\left(\sqrt{N_\phi}-1\right)^{-1}, 0^+\right)$ and $(\phi_4, \psi_4) \to (0^-, 1^-)$; $x^-$ and $x^+$ respectively denote values that are infinitesimally smaller or larger than $x$ (where the relevant $x$ values are 0 and 1). This indicates that the points $(\phi_1, \psi_1)$ and $(\phi_2, \psi_2)$ are within the physical region of first quadrant of the plane $(\phi, \psi)$, while $(\phi_3, \psi_3)$ and $(\phi_4, \psi_4)$, whose $\phi$ values are negative, are unphysical.

For values of the heterotypic interaction $\chi_{\phi\psi} < \chi_{\phi\psi}^{lc}$, we consider the limiting case $\chi_{\phi\psi} \to 0$ for which the expressions for the four points simplify, and use these simplified expressions to select the two physical points. However, we note that in contrast to the limit $\chi_{\phi\psi} \to \infty$, in the limit $\chi_{\phi\psi} \to 0$, (whose typical phase diagram is presented in Fig. 3B in the main text), the binodal does not form a closed loop; instead, its two branches intersect the line $\phi + \psi = 1$. Due to the relation of the binodal and spinodal, this suggests that the points along the spinodal corresponding to the minimal and maximal values $\psi$, are in the first quadrant $\phi, \psi > 0$, although not necessarily in the physical region of this quadrant, $\phi + \psi \leq 1$. In the limit $\chi_{\phi\psi} \to 0$, we find that the points $(\phi_1, \psi_1)$ and $(\phi_2, \psi_2)$ are in the first quadrant while $(\phi_3, \psi_3)$ and $(\phi_4, \psi_4)$ are not. Thus, the points $(\phi_1, \psi_1)$ and $(\phi_2, \psi_2)$ are the physical ones for values of $\chi_{\phi\psi} < \chi_{\phi\psi}^{lc}$ as well as for $\chi_{\phi\psi} > \chi_{\phi\psi}^{uc}$.

When the heterotypic interaction strength is equal to either of the critical ones, $\chi_{\phi\psi}^{lc}$ or $\chi_{\phi\psi}^{uc}$, the spinodal collapses to a single point where the two points of maximal and minimal $\psi$ are identical. Therefore, equating the expressions for either the $\phi$ value (Eqs. 67 and 69), or their corresponding $\psi$ values (Eqs. 59 and 61) that give the two extremal points along the spinodal, provides an equation whose solutions are the critical heterotypic interactions $\chi_{\phi\psi}^{lc}$ and $\chi_{\phi\psi}^{uc}$. The equation that results from equating either Eqs. 67 and 69, or Eqs. 59 and 61, is the same quartic equation of $\chi_{\phi\psi}$:

$$\left(1 + \frac{\chi_{\phi\phi}}{\chi_{\phi\psi}^2}\left(1 - \frac{1}{N_\psi}\right) - \frac{2}{\chi_{\phi\psi}}\left(1 + \frac{1}{\sqrt{N_\phi}}\right)\right)^2 - \frac{4}{\chi_{\phi\psi}^2}\frac{1}{N_\psi}\left(\left(1 + \frac{1}{\sqrt{N_\phi}}\right)^2 - \chi_{\phi\phi}\right) = 0$$

which has four roots. However, substituting two of these four roots into Eqs. 59 and 61 results in negative values of $\psi$, so that these two values $\chi_{\phi\psi}$ cannot be the critical ones, since the spinodal point must lie in the physical region of concentration space, $\phi, \psi \geq 0$ and $\phi + \psi \leq 1$. The two remaining values of $\chi_{\phi\psi}$ are thus the critical ones; as required by the definitions of $\chi_{\phi\psi}^{lc}$ and $\chi_{\phi\psi}^{uc}$, the smaller one is $\chi_{\phi\psi}^{lc}$ and the larger one is $\chi_{\phi\psi}^{uc}$, which are expressed as follows:

$$\chi_{\phi\psi}^{lc} = \left(1 - \frac{1}{\sqrt{N_\psi}}\right)\left(\left(1 + \frac{1}{\sqrt{N_\phi}}\right) - \sqrt{\left(1 + \frac{1}{\sqrt{N_\phi}}\right)^2 - \chi_{\phi\phi}}\right) \tag{75}$$

$$\chi_{\phi\psi}^{uc} = \left(1 + \frac{1}{\sqrt{N_\psi}}\right)\left(\left(1 + \frac{1}{\sqrt{N_\phi}}\right) + \sqrt{\left(1 + \frac{1}{\sqrt{N_\phi}}\right)^2 - \chi_{\phi\phi}}\right) \tag{76}$$

The two critical values for $\chi_{\phi\psi}$ that we have found, determine the nature of LLPS in ternary systems of this class. For $\chi_{\phi\psi} < \chi_{\phi\psi}^{lc}$, the LLPS is segregative, while for $\chi_{\phi\psi} > \chi_{\phi\psi}^{uc}$, the LLPS is associative; in between, when $\chi_{\phi\psi}^{uc} > \chi_{\phi\psi} > \chi_{\phi\psi}^{lc}$, the system does not exhibit LLPS and the mixed phase is the equilibrium state for all values of the concentrations $\phi$ and $\psi$.

###### 4.3.3 Class (3) - Small homotypic interaction: $\phi$ can only phase separate associatively for finite values of $\psi$

For ternary systems of this class, the homotypic interactions $\chi_{\phi\phi}$ are too small to facilitate segregative LLPS even when supplemented by crowding interactions. This indicates that, even if the multivalent binding partner is replaced with a crowder (i.e., one that has only excluded volume interactions with the other solute and the solvent, but no heterotypic interactions), the system will not phase separate. As explained in the subsection "Case I - Homotypic ($\chi_{\phi\phi} > 0, \chi_{\phi\psi} = 0$): Concentrated phase rich in one solute (segregative LLPS)", this means that the homotypic interaction strength is less than $\chi_{\phi\phi}^m$, which is specified in Eq. 28. Since heterotypic interactions inhibit segregative LLPS, systems of this class, which do not exhibit segregative LLPS for $\chi_{\phi\psi} = 0$ (i.e., where $\psi$ is a crowder), will not do so also in the presence of a positive heterotypic interaction $\chi_{\phi\psi} > 0$. However, such systems can still undergo associative LLPS if the heterotypic interaction $\chi_{\phi\psi}$ is large enough. Below, we determine the critical heterotypic interaction above which, class (3) systems undergo associative LLPS.

The critical value of the heterotypic interaction can be calculated from the spinodal curve. Since the spinodal and binodal must both exist when one does, the critical heterotypic interaction for which the spinodal appears is the one above which the system phase separates associatively. Moreover, since the Hessians of classes (2) and (3) are identical, the critical interaction parameters $\chi_{\phi\psi}^{lc}$ and $\chi_{\phi\psi}^{uc}$ respectively given by Eqs. 75 and 76, mark the disappearance and appearance of the spinodal for this class as well. The lower critical parameter, $\chi^{lc}_{\phi\psi}$, does not give rise to a physical transition for class (3) systems. This is because the concentrations of solutes, $\phi$ and $\psi$, at the spinodal point for $\chi_{\phi\psi} = \chi^{lc}_{\phi\psi}$, (obtained by substitution of $\chi^{lc}_{\phi\psi}$ from Eq. 75 into Eqs. 59 and 67), results in non-physical concentrations that satisfy $\phi + \psi > 1$ when $\chi_{\phi\phi} < \chi^{m}_{\phi\phi}$. In contrast, the upper critical interaction $\chi^{uc}_{\phi\psi}$, which marks the onset of associative LLPS in class (2) systems is physically relevant for class (3) systems and is the critical parameter above which the systems undergo associative LLPS. It is important to note that in the limit of vanishing homotypic interactions $\chi_{\phi\phi} \to 0$, the critical heterotypic interaction $\chi^{uc}_{\phi\psi}$, given by Eq. 76, coincides with the critical interaction $\chi^{c}_{\phi\psi}$, given by Eq. 43 for system of case II. This supports the consistency of our theory since for $\chi_{\phi\phi} \to 0$, the systems of case III are equivalent to those of case II. Furthermore, the definitions of $\chi^{c}_{\phi\psi}$ and $\chi^{uc}_{\phi\psi}$ imply that $\chi^{c}_{\phi\psi} > \chi^{uc}_{\phi\psi}$ for any $\chi_{\phi\phi} > 0$, which suggests that even weak homotypic interactions stabilize associative LLPS, driven by heterotypic interactions. This is consistent with the results (Fig. 5D) and explanations (see subsection "Heterotypic interactions: in the absence or together with homotypic ones") provided in the main text.

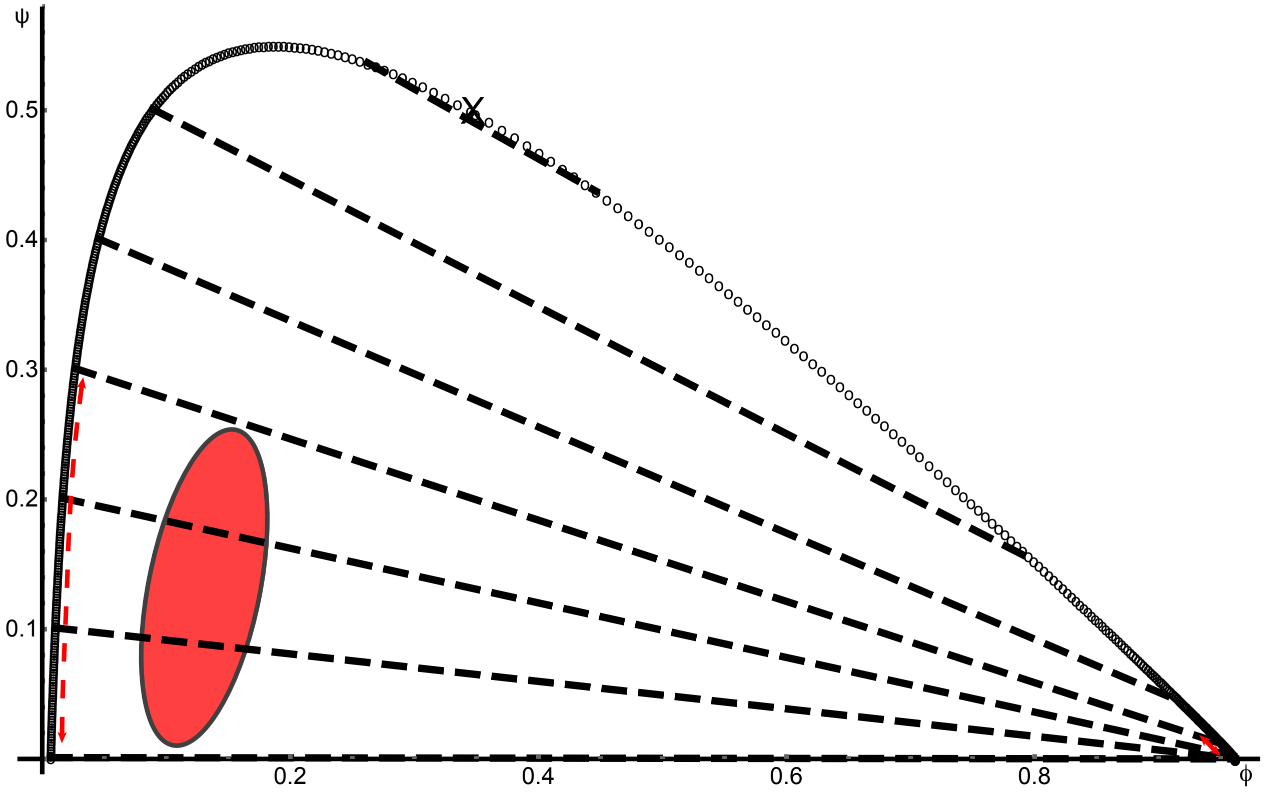


Figure 1: A ternary system that exhibits a phase diagram with highly splayed tie-lines. Binodal points are plotted as black, open circles, tie-lines as black, dashed lines, and the critical point with a black "X" mark. To highlight the possible concentration buffering capacities of such splayed phase diagram, an example of a noise-ellipse that is almost perpendicular to the tie-lines is plotted in red. The projected variations of the concentrations in the two coexisting phases due to the noise in the overall concentrations are indicated by red, dashed double headed arrows. Due to the large mismatch between the orientations of the noise-ellipse and the tie-lines, there is a wide range of possible equilibrium concentrations, so that conventional concentration buffering that buffers noise in both dilute and dense phases is not effective. However, as can be seen, the large splay of the tie-lines renders the dense phase robust to the large noise in the overall concentrations, while it augments the effect of the noise in the dilute phase. The parameters used to generate the phase diagram are $N_\phi = 2$, $N_\psi = 15$, $\chi_{\phi\phi} = 6$, and $\chi_{\phi\psi} \approx 2.91$.

| Critical interaction symbol | Full expression as a function of the relevant lengths and interactions |
|---|---|
| $\chi^{c}_{\phi\phi}$ | $\left(1+\frac{1}{\sqrt{N_\phi}}\right)^2$ |
| $\chi^{m}_{\phi\phi}$ | $\left(\frac{1}{\sqrt{N_\phi}}+\frac{1}{\sqrt{N_\psi}}\right)^2$ |
| $\chi^{c}_{\phi\psi}$ | $2\left(1+\frac{1}{\sqrt{N_\psi}}\right)\left(1+\frac{1}{\sqrt{N_\phi}}\right)$ |
| $\chi^{b}_{\phi\psi}$ | $\frac{1}{2}\chi_{\phi\phi}-\frac{1}{2}\left(\frac{1}{\sqrt{N_\phi}}+\frac{1}{\sqrt{N_\psi}}\right)^2$ |
| $\chi^{t}_{\phi\psi}$ | $\chi_{\phi\phi}/\left(1+\frac{1}{\sqrt{N_\phi}}\right)$ |
| $\chi^{lc}_{\phi\psi}$ | $\left(1-\frac{1}{\sqrt{N_\psi}}\right)\left(\left(1+\frac{1}{\sqrt{N_\phi}}\right)-\sqrt{\left(1+\frac{1}{\sqrt{N_\phi}}\right)^2-\chi_{\phi\phi}}\right)$ |
| $\chi^{uc}_{\phi\psi}$ | $\left(1+\frac{1}{\sqrt{N_\psi}}\right)\left(\left(1+\frac{1}{\sqrt{N_\phi}}\right)+\sqrt{\left(1+\frac{1}{\sqrt{N_\phi}}\right)^2-\chi_{\phi\phi}}\right)$ |

Table 1: Explicit, analytical expressions of the critical interactions energies derived in “Supplementary results”

| Topological property | lower bound on $\chi_{\phi\phi}$ value | upper bound on $\chi_{\phi\phi}$ value | lower bound on $\chi_{\phi\psi}$ value | upper bound on $\chi_{\phi\psi}$ value |
|---|---|---|---|---|
| binodal intersects the line $\psi = 0$ | $\chi^{c}_{\phi\phi}$ | $\infty$ | Independent | Independent |
| binodal intersects the line $\phi + \psi = 1$ | $\chi^{m}_{\phi\phi}$ | $\infty$ | 0 | $\chi^{b}_{\phi\psi}$ |
| Segregative LLPS: Continuous transition | $\chi^{c}_{\phi\phi}$ | $\infty$ | 0 | $\chi^{t}_{\phi\psi}$ |
| Associative LLPS: Continuous transition | $\chi^{c}_{\phi\phi}$ | $\infty$ | $\chi^{t}_{\phi\psi}$ | $\infty$ |
| Segregative LLPS: Re-entrant transition | $\chi^{m}_{\phi\phi}$ | $\chi^{c}_{\phi\phi}$ | 0 | $\chi^{lc}_{\phi\psi}$ |
| Associative LLPS: Re-entrant transition | 0 | $\chi^{c}_{\phi\phi}$ | $\chi^{uc}_{\phi\psi}$ | $\infty$ |

Table 2: Classification of phase-diagram topologies: For each topological property considered in the section “Supplementary results”, we specify the ranges of the homotypic and heterotypic interactions (lower and upper bounds) for which the given property applies. Motivated by the absence of specific repulsive interactions (above and beyond excluded volume, which for a screening length of 1nm also subsumes the Coulomb repulsion of like charges moieties) in biological systems, we limit all interactions to be attractive (positive parameters). The terms “Continuous” and “Re-entrant” describe two possible types of transitions between segregative and associative LLPS, as we have analyzed here. Intersection of the binodal with the line $\psi = 0$ is independent of the heterotypic interaction because in the absence of solute $\psi$, the heterotypic interactions have no effect.

[3] Samuel Safran. *Statistical thermodynamics of surfaces, interfaces, and membranes.* CRC Press, 2018.

[4] Miroslav Fiedler. *Matrices and graphs in geometry.* Number 139. Cambridge University Press, 2011.

[5] Dan Deviri and Sam A Safran. Equilibrium size distribution and phase separation of multivalent, molecular assemblies in dilute solution. *Soft Matter*, 2020.

[6] Pierre-Gilles De Gennes and Pierre-Gilles Gennes. *Scaling concepts in polymer physics.* Cornell university press, 1979.

[7] Kurt Binder and Peter Fratzl. Spinodal decomposition. *Phase transformations in materials*, pages 409–480, 2001.

[8] unpublished.
